# MicroDUET: a dual-headed genomic foundation model for pathogen surveillance

**DOI:** 10.64898/2026.11.11.765853

**Authors:** Michael Barber, Carl J. E. Suster, Jingyi Lei, Marya Getchell, Suci Wulandari, Vitali Sintchenko, Paul M. Pronyk

## Abstract

Pathogen genomic surveillance works from raw reads, assembled genomes and alignments, representations that are slow and costly to compare at scale. Compact embeddings from genomic language models offer an alternative, but surveillance requires resolving both evolutionary ancestry (phylogeny) and phenotypic capability (function) - which need not co-vary. Here we present MicroDUET, which adapts a 26.6-million-parameter ProkBERT backbone with two 3.5-million-parameter adapter heads: a phylogeny head trained contrastively on genome windows from the same microbes, and a function head trained to identify genes across taxa. On held-out retrieval benchmarks, the phylogeny head returned the correct species for 80.3% and 66.6% of cellular and viral queries, and the function head the correct gene family for 97.3% and 97.2%, reaching 74.4% accuracy on an independent antimicrobial resistance-class benchmark. Across all three benchmarks MicroDUET outranked 17 other genomic encoders. Without training on lineage, clade or drug-susceptibility tasks, MicroDUET identified *Mycobacterium tuberculosis* and mpox virus lineages and clades (0.965, 0.975 accuracy), discriminated rifampicin resistance without lineage adjustment (AUROC 0.985), and retrospectively identified an emerging mpox lineage as anomalous (AUROC 0.974). MicroDUET operates 10^3^ - 10^4^ times faster than single-threaded sequence-based alignments. MicroDUET therefore provides a compact, reusable representation for scalable pathogen surveillance and sequence triage.

## Introduction

Pathogen genomic surveillance uses sequence variation to identify circulating lineages, detect emerging variants, investigate transmission, and infer clinically relevant phenotypes such as antimicrobial resistance. Genomic epidemiology has consequently become central to international strategies for epidemic and pandemic-preparedness^1–3^. However, translating genomic data into actionable intelligence remains dependent on pathogen-specific pipelines involving reference alignment, variant calling, quality filtering, phylogenetic reconstruction, and specialist interpretation. Although essential, these workflows can be time consuming and require substantial computational resources and technical expertise. These demands are particularly relevant in resource-constrained settings, where limited bioinformatics capacity and computing infrastructure must support surveillance tasks across multiple priority pathogens,^4^ contributing to persistent global inequalities in genomic surveillance capacity.^5^

Genomic language models and sequence embeddings have emerged as promising computational approaches across human functional genomics, microbial biology, and nucleotide sequence classification.^6–10^ By learning compact representations of nucleotide sequences, these models can encode reads, genes, contigs, or whole genomes as numerical vectors. These embeddings capture biological signals at multiple scales, from taxonomic classification and gene identity to structural and functional variation.^8,11^ In pathogen genomics, compact language models such as ProkBERT have demonstrated the potential of microbial pretraining to capture promoter architecture, phage insertion boundaries, and taxonomic divergence.^11^ More specialized models, including DNABERT-S, Scorpio, and vir2vec, have demonstrated capabilities in species discrimination, evolutionary relatedness, and antimicrobial resistance class identification.^6,9,12–14^

Genomic surveillance asks two distinct questions of the same sequence: how an isolate relates to others in circulation, and what that isolate is capable of. These questions can favour different relationships in embedding space.^15^ Sequences with shared ancestry may cluster together despite differences in resistance phenotype,^11,13^ whereas similar functional determinants can occur across divergent genetic backgrounds.^15,16^ Optimising a representation for one relationship may therefore reduce performance on the other. This motivates testing whether separate adaptations of a compact shared encoder can support evolutionary and functional inference while retaining information useful across surveillance tasks.

A second barrier is computational. Large genomic foundation models, including Evo 2 and the 2.5-billion-parameter Nucleotide Transformer impose substantial inference costs that can constrain routine analysis on standard laboratory hardware.^7,8^ This can limit their use in surveillance programmes, particularly in resource-limited settings. Whether compact genomic language models can retain sufficiently rich evolutionary and functional information for diverse public health surveillance tasks while remaining deployable on widely available hardware remains an important question.

*Mycobacterium tuberculosis* (MTB) and mpox virus (MPXV) provide representative examples of priority pathogens for public health surveillance and contrasting archetypes for evaluating these capabilities. *M. tuberculosis* is a predominantly clonal, slowly evolving bacterial pathogen in which single-nucleotide variants can confer clinically relevant antimicrobial resistance.^17^ MPXV, by contrast, is a large double-stranded DNA virus whose recent international epidemics have involved distinct viral clades and lineages, structural genomic variation, and characteristic mutational signatures associated with host APOBEC3 cytidine deaminase activity.^18,19^ Together, these pathogens allow us to evaluate complementary evolutionary and functional representations across substitution-driven variation, population structure, resistance and gene-scale loss.

Here, we introduce MicroDUET (Microbial-focused Dual-headed Unified Embeddings Transformer, Fig. 1), designed to provide complementary evolutionary and functional representations, support CPU-only deployment, generalise to surveillance tasks absent from training, and enable an index-once, query-many workflow. With a frozen ProkBERT-mini-long backbone^11^ (26.6 million parameters) selected for predictive performance and throughput among compact encoders, we trained two lightweight 3.5-million-parameter transformer adapter heads: a phylogeny head via contrastive whole-genome learning, adapting the objective introduced by DNABERT-S^13^, and a function head via supervised gene family classification across divergent taxa. Both heads were trained on these two objectives alone; no lineage, clade, drug-susceptibility or mutational-signature labels were used at any training stage.

**Figure 1.**
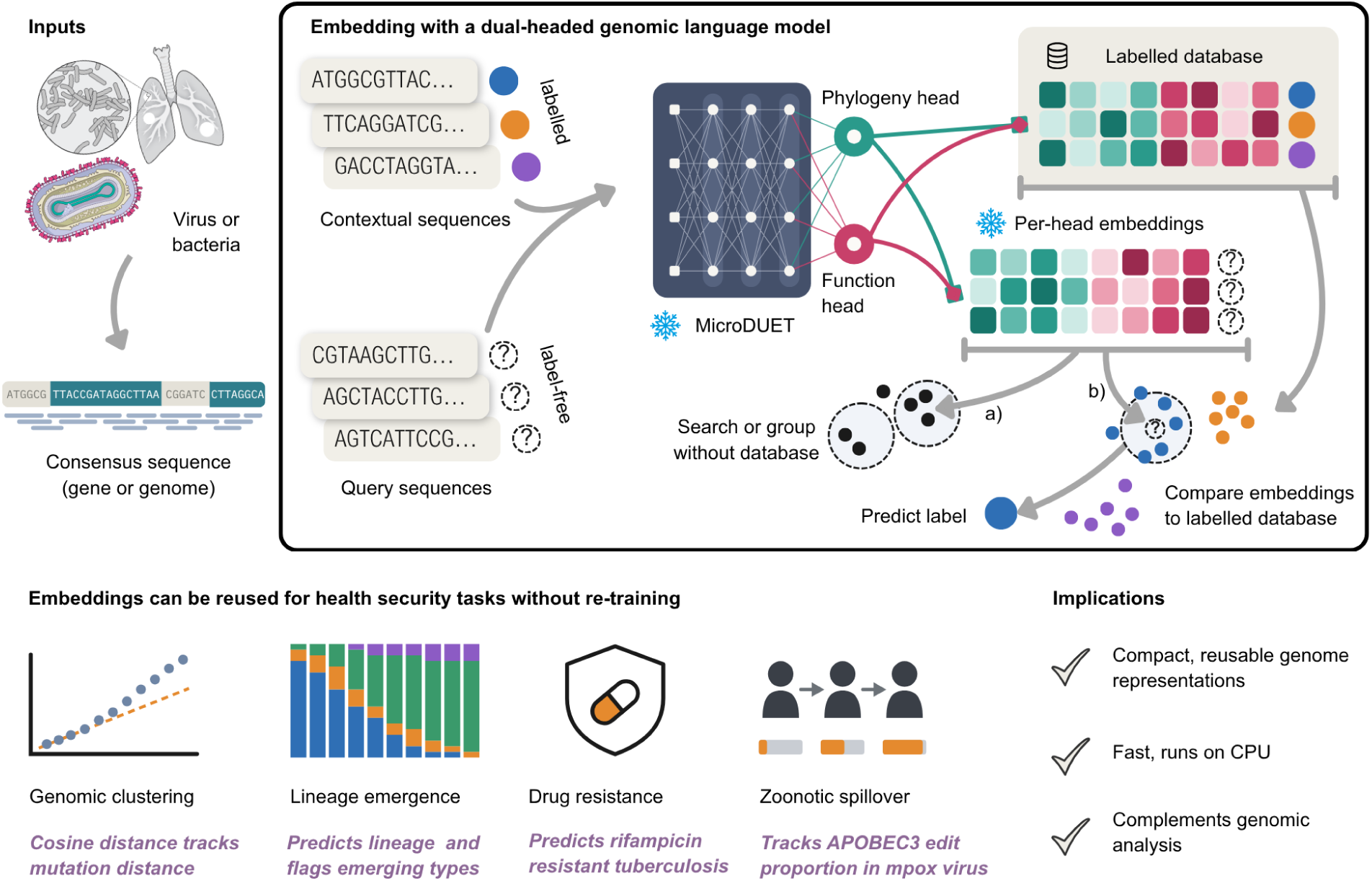
MicroDUET schematic. MicroDUET is a two-headed genomic language model that produces compact pathogen representations for surveillance tasks. DNA sequences from bacteria or viruses are embedded by either head into fixed length vectors (green and red squares) depending on the task required. Query sequences (black filled or dashed circles) are embedded and a) assessed without labels for similarity and divergence - such as identifying emergent strains or b) compared against a labelled database of embeddings linked to known characteristics (coloured circles) to predict the genotype or phenotype of the query sequence. The same embeddings can be reused across public health tasks, including inference of population structure, lineage novelty and drug resistance, without task-specific retraining. Snowflakes denote frozen components.

We then evaluated MicroDUET zero-shot on *M. tuberculosis* and MPXV sequences, assessing mutational distance, mutation consequence, lineage and clade assignment, rifampicin resistance discrimination, emerging variant detection, and APOBEC3-associated mutational patterns, with model weights remaining fixed during these evaluation tasks. We demonstrate that MicroDUET resolves lineage, clade and resistance tasks that were absent from both training objectives, while stored embeddings accelerate routine surveillance operations 10^3^-10^4^ fold faster than conventional alignment approaches on standard workstation hardware. By enabling rapid genomic analysis, this lightweight approach could help accelerate the detection of emerging threats and help surveillance programmes rapidly prioritise sequence analyses in settings with limited resources.

## Results

### A shared compact backbone supports complementary evolutionary and functional representations

Under a CPU-only deployment constraint, we screened 17 pretrained genomic encoders on three measures (Fig. 2a; Supplementary Data Table 1). For each query we took its nearest neighbour by embedding cosine similarity and asked whether it shared the query’s label. In species identification, the query was a genome window of 100 bp–10 kb (1,000 cellular and 920 viral windows, each set from 200 species), and a positive hit was a neighbour from the same species. In gene-function retrieval, the query was a coding sequence (1,200 cellular and 1,000 viral), and a hit was a neighbour with the same annotated gene name. Neighbours from the same genus or virus taxon were excluded to reduce retrieval driven by shared taxonomic background (see methods). Throughput was measured as megabases embedded per minute. DNABERT-S led cellular species retrieval (77.0%) and ProkBERT-mini-long led viral species (61.7%) and viral gene-function retrieval (83.9%), while the fastest CPU encoder, PathoLM (1.26 megabases per minute), matched species for only 17.7% of cellular neighbours. Larger parameter counts did not consistently improve retrieval in this screen. Evo2-7B and NT-2.5B^7,8^ achieved cellular species-retrieval accuracies of 24.0% and 37.0%, respectively, compared with 57.9% for ProkBERT-mini-long^11^. Their inference throughput was substantially lower; NT-2.5B, for example, embedded 0.005 megabases per minute on CPU, compared with 0.409 for ProkBERT-mini-long (Supplementary Data Table 1). Because throughput and retrieval accuracy did not rank together, we retained ProkBERT-mini-long as a frozen backbone for further adaptation to public health tasks.

**Figure 2.**
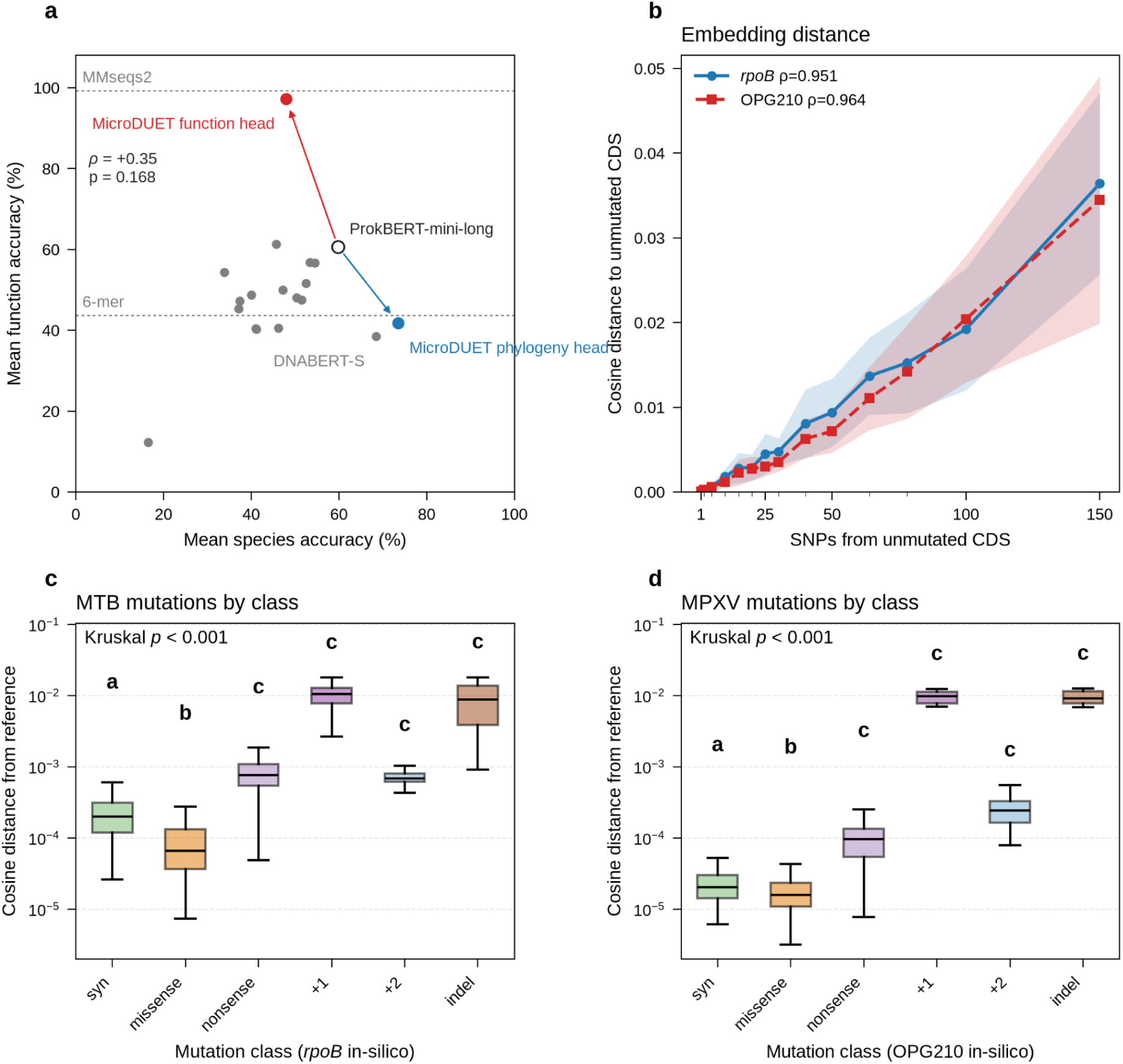
MicroDUET’s two heads specialise in opposite directions, and embedding distance tracks substitution count and differs among variant classes. **a**, Mean species accuracy (mean of cellular and viral species retrieval, k = 1) against mean gene-function accuracy (mean of cellular and viral gene-function retrieval) for ProkBERT-mini-long (open circle), the MicroDUET phylogeny head (blue) and function head (red), and 16 other pretrained genomic language models (grey). Arrows show the shift from the ProkBERT backbone. Dashed lines show the mean function accuracy of MMseqs2 translated alignment and of a six-mer composition baseline. We did not detect a correlation between species and function accuracy across the 17 pretrained models (Spearman ρ = 0.35, *P* = 0.168). **b**, Pairwise cosine embedding distance as a function of accumulated nucleotide substitutions across 30 independent mutational paths (1 to 150 single-nucleotide polymorphisms, SNPs) in *M. tuberculosis rpoB* gene (3,519 bp) and mpox virus (MPXV) OPG210 gene (5,643 bp) using the MicroDUET phylogeny head (n = 420 pairs per gene; *rpoB* Spearman ρ = 0.951, replicate-bootstrap 95% CI 0.943–0.959; OPG210 ρ = 0.964, 0.956–0.971). **c, d**, Cosine distance from the unmutated reference for in-silico variants of *rpoB* (c, *n* = 4,626) and OPG210 (d, *n* = 2,988). Substitutions are every GC→AT change. Frameshifts and in-frame indels were drawn as 50 each. Distances differed among the six classes (Kruskal–Wallis *H* = 1,810, *p* < 0.001 for *rpoB*; *H* = 678, *p* < 0.001 for OPG210). Letters mark classes not separated by Holm-adjusted Dunn tests on the pooled ranks. Shared letters do not imply equal medians. Boxplots show the first to third quartiles, median, and 1.5x interquartile range.

With the backbone frozen, contrastive training of the MicroDUET phylogeny head increased same-species retrieval to 80.3% for cellular and 66.6% for viral sequences, versus 57.9% and 61.7% for unadapted ProkBERT (Fig. 2a; Supplementary Data Table 1). Gene-family supervision of the function head instead raised gene-function retrieval to 97.3% cellular and 97.2% viral, while cellular species retrieval fell to 31.6% (Supplementary Data Table 2). As an independent function-oriented check, we also scored retrieval of resistance class across the NCBI AMRFinderPlus reference gene catalogue,^20^ a curated set of antimicrobial resistance genes used to detect resistance determinants in clinical isolates; here the labels are resistance classes rather than the gene families the head was trained to separate. Across 398 coding sequences spanning 23 resistance classes, the MicroDUET function head recovered the correct class for 74.4% of queries, versus 56.3% for the unadapted backbone (Supplementary Data Table 1). The two heads therefore specialise on complementary retrieval objectives while sharing a single pretrained encoder.

### Phylogeny-head embedding distance tracks substitution count and mutation consequence

To determine whether the continuous geometry of the phylogeny head preserves discrete mutational separation, we simulated 30 independent nested mutational trajectories across 14 substitution counts (1 to 150 single-nucleotide polymorphisms, SNPs) in two loci: *M. tuberculosis rpoB* gene (3,519 bp), which carries the mutations confirming resistance to the first-line antituberculous drug rifampicin,^16^ and MPXV OPG210 (5,643 bp), a surface membrane glycoprotein that suppresses T-cell activation.^21^ Pairwise cosine distance tracked substitution count in both loci (Spearman ρ = 0.951 for *rpoB*, replicate-bootstrap 95% CI 0.943–0.959; ρ = 0.964 for OPG210, 95% CI 0.956–0.971; *n* = 420 pairs per locus; Fig. 2b). In the low-divergence regime relevant to transmission investigation (≤ 15 SNPs) the relationship was approximately linear, with per-SNP displacement scaling inversely with locus length; dispersion across replicates widened with divergence, so individual pairwise distances should not be converted to SNP equivalents.

We next asked whether displacement reflects mutation consequence as well as count. Across 4,626 and 2,988 in silico coding variants in *rpoB* and OPG210 genes respectively, displacement differed by consequence class (Fig. 2c,d). Frameshift and indel variants displaced vectors furthest from the unmutated reference. In *rpoB*, the median cosine distance rose from synonymous substitutions (2.0 × 10⁻⁴) to nonsense mutations (7.68 × 10⁻⁴), in-frame indels (8.82 × 10⁻³) and +1 frameshifts (1.05 × 10⁻²). The same sensitivity to disrupted coding structure held for deletions carried by circulating genomes: four published MPXV events^18,22,23^ lay 15× to 1,800× the background median in whole-genome cosine distance to the nearest intact neighbour (Supplementary Data Fig. 1). Phylogeny-head geometry therefore encodes substitutions, whether a variant disrupts coding structure, and gene-scale loss at genome scale, although in silico variants and reference-relative distances do not establish the phenotypic consequence of any individual change.

### Phylogeny-head neighbourhoods recover lineage, clade and epidemic composition without task-specific labels

To evaluate whether the phylogeny head resolves population structure in clinical isolates, we examined 3,638 unique sequences across 39 antimicrobial resistance-associated loci^16^ from 1,766 clinical *M. tuberculosis* isolates, alongside *rpoB* (*n* = 347) and OPG210 (*n* = 404) sequences for lineage and clade structure. In two-dimensional projection using Uniform Manifold Approximation and Projection (UMAP)^24^, the 39 loci formed distinct, non-overlapping clusters corresponding to individual genes without sequence alignment or prior annotation (Fig. 3a). Within individual loci, nearest-neighbour geometry captured broad-scale population structure: leave-one-out nearest neighbours (k = 5, inverse cosine weighting) on *rpoB* matched the isolate’s major lineage with a mean same-lineage weight (the distance-weighted fraction of an isolate’s neighbours carrying the same lineage) of 0.943 (95% CI 0.925–0.960; chance 0.333) and a balanced accuracy of 0.965 (95% CI 0.942–0.984; Fig. 3b; Supplementary Data Fig. 2a). In MPXV, OPG210 neighbourhoods assigned clinical isolates to their WHO clade^25^ with a mean same-clade weight of 0.982 (95% CI 0.971–0.991; chance = 0.250; Fig. 3c), yielding median weights of 1.00 for clades Ia (n = 63), Ib (n = 21), and IIb (n = 315), and 0.931 for clade IIa (n = 5). In leave-one-out classification, OPG210 achieved a balanced accuracy of 0.975 (95% CI 0.941–1.000), correctly classifying all five clade IIa isolates, again above the unadapted backbone (Supplementary Data Fig. 2b; Supplementary Data Table 2). A single gene can resolve clades because lineage-specific SNPs and small structural mutations are distributed across much of the MPXV genome^26^.

**Figure 3.**
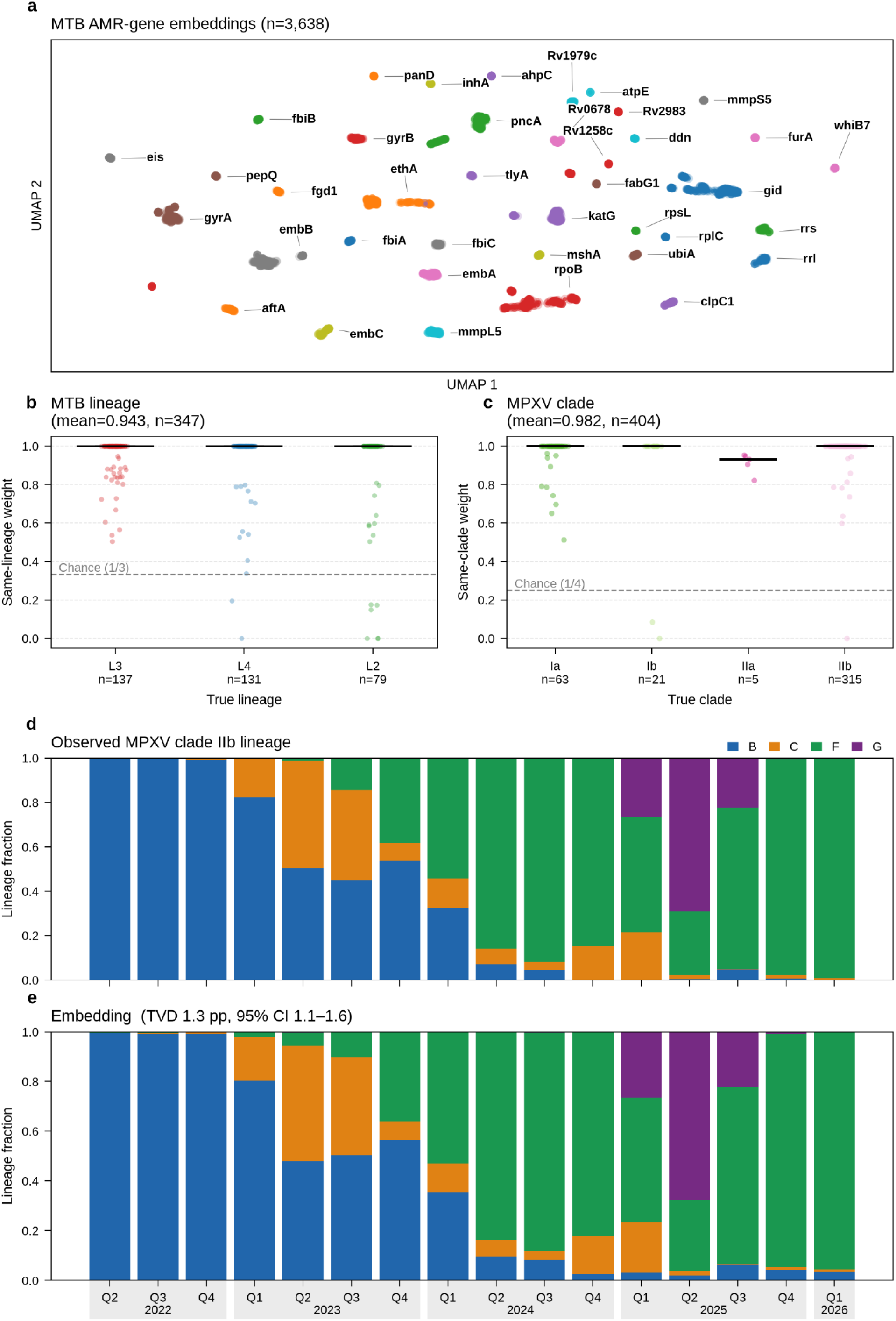
MicroDUET phylogeny head resolves population structure and tracks epidemic lineage dynamics. **a**, Two-dimensional UMAP projection of 3,638 unique coding sequences across 39 antimicrobial resistance-associated loci from 1,766 clinical *M. tuberculosis* isolates, coloured by locus. Sequences partition into discrete gene clusters without alignment or sequence annotation. **b,** Leave-one-out nearest-neighbour same-lineage weight (k = 5, inverse cosine weighting) for *rpoB* sequences across 347 clinical *M. tuberculosis* isolates spanning lineages 2, 3, and 4 (mean weight = 0.943, 95% CI 0.925–0.960; dashed line indicates analytical chance expectation of 0.333). **c**, Nearest-neighbour same-clade weight (k = 5) for OPG210 sequences across 404 clinical MPXV isolates across WHO clades Ia (n = 63), Ib (n = 21), IIa (n = 5), and IIb (n = 315; mean weight = 0.982, 95% CI 0.971–0.991; dashed line indicates the equal-class reference of 0.250). **d, e,** Longitudinal reconstruction of circulating lineage dynamics across 8,738 whole-genome MPXV clade IIb consensus genomes collected over 16 calendar quarters (2022-Q2 to 2026-Q1). Observed lineage proportions (d) compared with out-of-fold nearest-neighbour lineage predictions (e) across lineages B, C, F, and G (total variation distance 1.3 percentage points, 95% CI 1.1–1.6). Box plots: center lines indicate the median; box limits indicate the first and third quartiles; whiskers extend to 1.5× the interquartile range.

For structure within a clade this was not the case: most alleles of any individual gene are shared between lineages, and no single-gene embedding approached the accuracy of whole-genome embeddings (Supplementary Data Fig. 3). We therefore evaluated whole-genome embeddings across 8,738 clade IIb consensus genomes collected over 16 calendar quarters (2022-Q2 to 2026-Q1),^27^ 13 of which postdate the 6 January 2023 retrieval of the backbone pretraining corpus.^11^ Out-of-fold nearest-neighbour classification (*k* = 5) reconstructed the observed quarterly succession of circulating lineages B, C, F and G (total variation distance 1.3 percentage points, 95% CI 1.1–1.6; Fig. 3d,e; Supplementary Data Fig. 4; Supplementary Data Fig. 2d), capturing the expansion of lineage B through 2022 and its progressive replacement by lineages C, F and G across 2024 and 2025. These results demonstrate that the phylogeny head captures both deep divergence between clades in single genes and cumulative genome-wide divergence during an evolving epidemic, with no lineage or clade labels at any stage of training.

### Function-head embeddings reduce lineage confounding in rifampicin resistance prediction

To evaluate whether sequence embeddings detect functional resistance without confounding by population structure,^15^ we scored coding sequences of the *rpoB* gene across 347 clinical *M. tuberculosis* isolates from the CRyPTIC Compendium^28,29^ with matched phenotypic drug-susceptibility testing for the first-line antitubercular rifampicin (pDST; 298 resistant, 49 susceptible phenotypes). On the unadapted ProkBERT backbone, nearest-neighbour retrieval reflected lineage and resistance together: leave-one-out classification (*k* = 5) assigned isolates to their major lineage with a balanced accuracy of 96.6% (95% CI 94.4–98.5%; Fig. 4a), and naive resistance prediction yielded AUROC 0.876 with 41 errors at *k* = 5 (33 false positives, 8 false negatives; Supplementary Data Table 2, Fig. 4b). Subtracting a lineage-average vector from each embedding increased resistance discrimination, raising AUROC to 0.943 (paired DeLong Δ = +0.068, *p* < 0.001; Fig. 4c).

**Figure 4.**
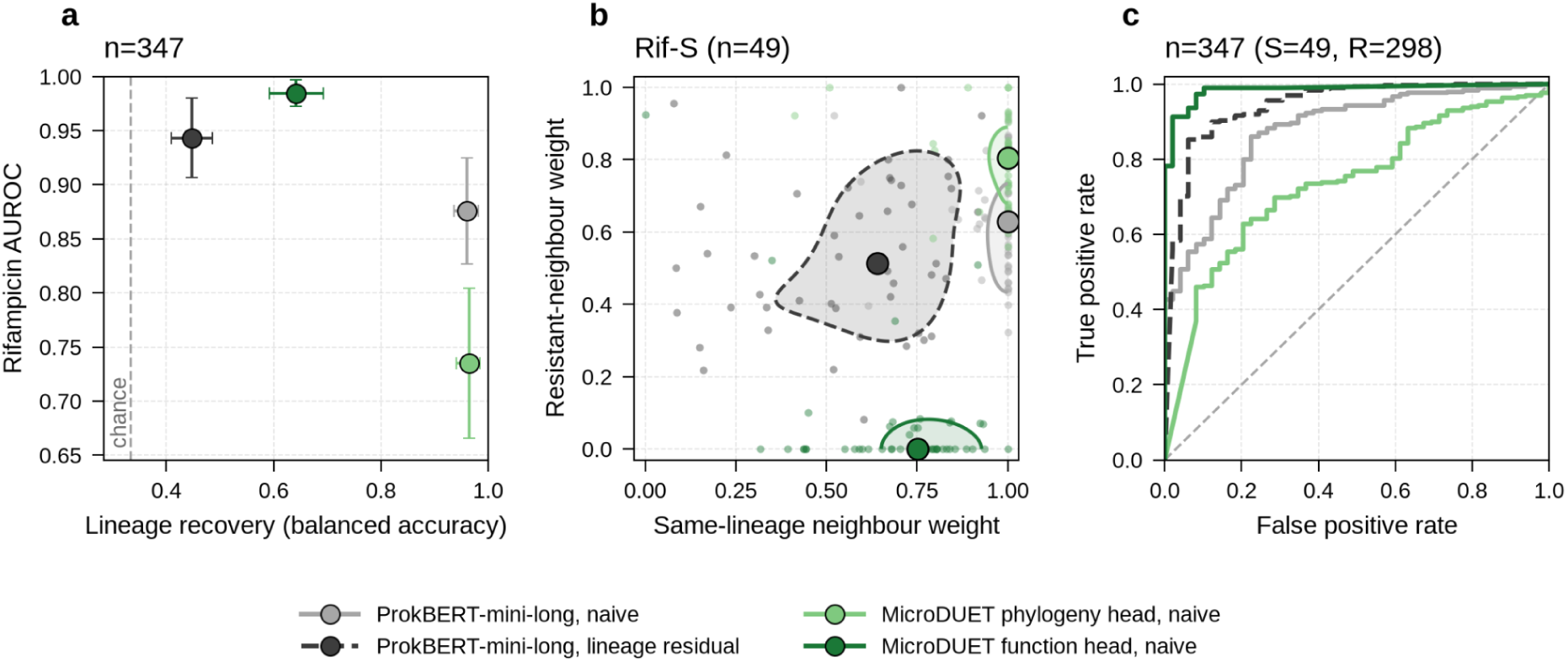
The function head discriminates rifampicin resistance without lineage adjustments. **a**, Balanced lineage accuracy versus rifampicin resistance prediction AUROC for 347 unique *rpoB* sequences (298 resistant, 49 susceptible) (out-of-fold, cosine distance-weighted). Error bars are 95% confidence intervals (2,000-resample percentile bootstrap; DeLong). The dashed line is chance balanced accuracy for three lineages (1/3). **b**, Sequences from rifampicin susceptible (Rif-S) cases (*n* = 49): neighbour weight on the sequence’s own lineage versus neighbour weight on rifampicin resistance. Contours are 50% highest-density regions; outlined points are coordinate-wise medians. **c**, Rifampicin resistance ROC curves for the same scores. The dashed line is chance. ProkBERT is shown as naive, and lineage-subtracted embeddings, MicroDUET is shown as phylogeny and function heads. DeLong tests, Holm-adjusted across six comparisons (*n* = 347): function versus species head, Δ = 0.249, *p* < 0.001; function head versus ProkBERT lineage-residual, Δ = 0.041, p = 0.035

The two adapter heads address this representation trade-off by specialising their objectives. On the phylogeny head, naive resistance discrimination fell to AUROC 0.735 (Fig. 4a, c) with 52 errors at *k* = 5 (45 false positives, 7 false negatives), consistent with neighbourhoods organised by lineage background. The function head discriminated resistance directly, at AUROC 0.985 with 22 errors (4 false positives, 18 false negatives; Supplementary Data Table 2), while lineage prediction accuracy fell to 68.4% (95% CI 63.7–73.3%; chance 33.3%; Fig. 4a). Lineage-mean subtraction, which improved the unadapted backbone, had little effect on the function head (0.985 naive versus 0.984 adjusted; Fig. 4c). Gene-family supervision thus reduced phylogenetic confounding and removed the need for post-hoc lineage adjustment in this cohort. Extending this to the rest of the CRyPTIC panel, neighbourhoods of gene embeddings using the function head separated resistant from susceptible isolates for nine further antitubercular drugs (Supplementary Data Fig. 8-9). For each drug, we chose one gene to embed based on reported resistance-associated mutations and compared to the genomic drug susceptibility call by TBProfiler using the WHO mutation catalogue. Discrimination ranged from AUROC 0.807 for pyrazinamide (*pncA*) to AUROC 0.954 for the fluoroquinolones (*gyrA*).

### Whole-genome embeddings identify unindexed lineages and recover APOBEC3-associated patterns

To assess whether stored embeddings detect emergent variants during an outbreak without retraining or taxonomic curation, we evaluated MicroDUET against sequences from the 2025 mpox epidemic in West Africa. Historical surveillance databases from the 2022 global epidemic are dominated by lineage B.1, which circulated primarily across Europe and the Americas. In contrast, the 2025 epidemic in Sierra Leone was driven by lineage G.1,^18^ a distinct clade IIb branch that diversified cryptically in West Africa before causing sustained local transmission. Querying 266 whole-genome G.1 consensus sequences against a static historical gallery of 3,842 clade IIb genomes from 2022 B.1, with 266 held-out 2022 B.1 genomes as in-distribution controls, cosine distance separated G.1 with an AUROC of 0.981 (95% CI 0.967–0.996; DeLong *p* < 0.001; Fig. 5a). In-distribution B.1 isolates clustered tightly against the gallery at a median cosine distance of 1.40 × 10⁻⁷ (interquartile range, IQR 7.19 × 10⁻⁸ to 2.54 × 10⁻⁷), whereas G.1 genomes showed a 166-fold displacement to a median of 2.33 × 10⁻⁵ (IQR 2.27 × 10⁻⁵ to 2.56 × 10⁻⁵). This displacement indicates that G.1 sequences occupy a region of embedding space unmapped by the 2022 epidemic index, reflecting accumulated branch-specific mutations that separate this lineage from the indexed B.1 background.

**Figure 5.**
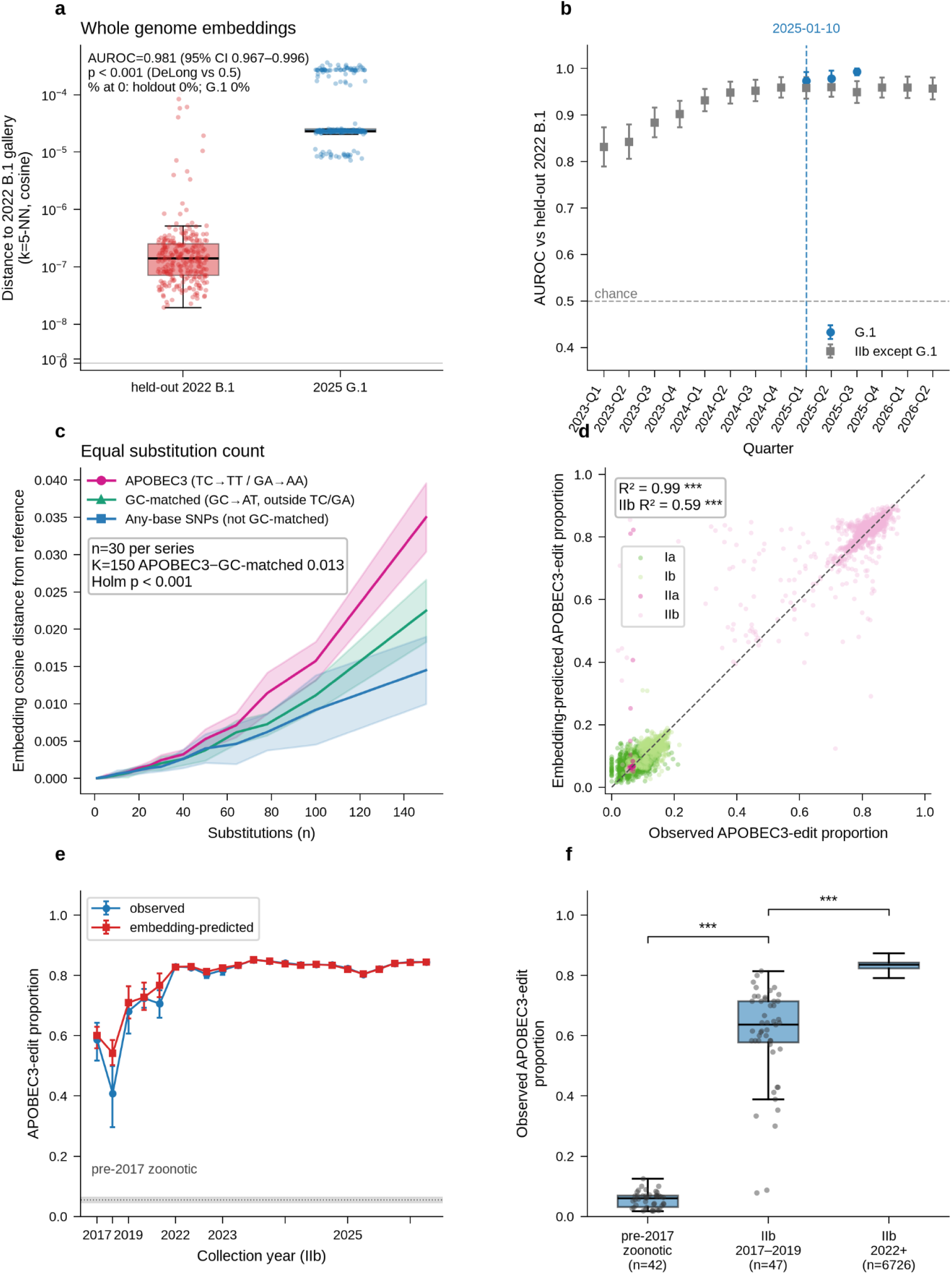
Gallery distance separates 2025 G.1 from 2022 B.1, and embeddings follow APOBEC3 editing. **a**, Fifth-nearest-neighbour cosine distance to clade IIb B.1 genomes collected in or before 2022 (n = 3,842), for held-out 2022 B.1 (n = 266) and G.1 (n = 266). AUROC = 0.981 (95% CI 0.967–0.996; DeLong, p < 0.001). **b**, Quarterly AUROC versus that holdout for G.1 (blue) and other IIb (grey); error bars, 95% CIs. blue dashed line is the earliest G.1 collection date (2025-01-10). (AUROC = 0.974 for G.1 detection in 2025-Q1-95% CI 0.955–0.992; n = 19; same test, p < 0.001). **c**, OPG210 cosine distance versus substitution count for APOBEC3 (TC→TT/GA→AA), GC-matched (GC→AT outside TC/GA) and other any-base edits (n = 30; band, ±1 s.d.). At 150 substitutions, Δ (APOBEC3 − GC-matched) = 0.0125 (Mann–Whitney test, Holm-adjusted over 14 counts, p < 0.001). **d**, Embedding-predicted versus observed APOBEC3-edit proportion for 8,563 unique genomes, coloured by clade; dashed line, y = x. R² = 0.985 (p < 0.001). **e**, Quarterly observed and embedding-predicted proportion, clade IIb (95% CIs). Grey band, pre-2017 baseline (year < 2017). **f**, Observed proportion: pre-2017 zoonotic (n = 42), IIb 2017–2019 (n = 47), IIb 2022+ (n = 6,726). One-sided Mann–Whitney test, later era higher (p < 0.001). In box plots, the box extends from the first quartile to the third quartile, with a line marking the median; whiskers span to the most extreme points within 1.5× the interquartile range.

Retrospective quarterly analysis separated G.1 from held-out 2022 B.1 controls in its first sampled quarter, 2025-Q1 (AUROC 0.974, 95% CI 0.955–0.992; n = 19), with AUROC increasing to 0.979 in 2025-Q2 and 0.993 in 2025-Q3 (Fig. 5b). Across 1,392 circulating non-G.1 clade IIb genomes collected between 2023-Q1 and 2024-Q4, lineages B, C, F and G represent continuous, clock-like drift along the original B.1 trunk; their median gallery distances rose gradually from 4.37 × 10⁻⁷ in 2023-Q1 to 1.71 × 10⁻⁶ in 2024-Q4, and their AUROC from 0.831 to 0.959 (Fig. 5b). The separation of G.1 instead reflects an abrupt branch shift from an unindexed West African A.2.2 background.^18^ Distance from a static index therefore flagged G.1 on first sampling, whereas ordinary drift along the B.1 trunk accumulated slowly.

Because MPXV evolution during circulation in humans is driven predominantly by host APOBEC3 cytidine deaminases rather than viral replication errors,^22,30^ we investigated whether MicroDUET captures this specific mutational mechanism. In controlled simulations holding total substitution count constant (n = 30 series up to 150 substitutions), canonical APOBEC3 dinucleotide transitions (TC→TT and GA→AA) displaced the phylogeny-head vector further than equal numbers of GC-matched substitutions outside those motifs (Δ = 0.0125 at 150 substitutions; Mann–Whitney, Holm-adjusted p < 0.001, Fig. 5c; Supplementary Data Fig. 5). Across 8,563 clinical MPXV consensus genomes, embedding distance predicted the genomic proportion of APOBEC3 mutations across clades (R² = 0.985, p < 0.001; Fig. 5d). Embedding trajectories accurately reconstructed the historical transition from pre-2017 zoonotic reservoirs into sustained human circulation in 2017-Q4 (Fig. 5e). Observed APOBEC3 proportions rose from a zoonotic median of 0.057 (n = 42) to 0.636 during the 2017–2019 Nigerian outbreak (n = 47, Mann–Whitney p < 0.001) and 0.833 across the 2022 global epidemic (n = 6,726, p < 0.001; Fig. 5f).^22,30,31^ MicroDUET thus captures asymmetric host-editing pressure and distinguishes sustained human transmission from zoonotic spillover in MPXV.

### The two adapter heads specialise on complementary tasks

Across the full benchmark panel (Supplementary Data Table 2), the two MicroDUET heads pull the representation in opposite directions. The phylogeny head was the stronger head for species, lineage and clade assignment, whereas the function head was the stronger head for gene-function retrieval, AMRFinder screening and rifampicin resistance. Each specialisation came at a measurable cost on the other head’s tasks: the function head’s lineage neighbour weight fell to 0.604 (phylogeny head 0.943), and the phylogeny head made n=45 false-positive rifampicin calls at k = 5 (function head n=4). Tasks driven by overall sequence divergence, including embedding–SNP correlation, G.1 detection, quarterly lineage-mix reconstruction and APOBEC3 edit-fraction prediction, were similar across heads (differences ≤ 0.016). Neither specialisation reduced sensitivity to overall sequence divergence, which both heads retain (Supplementary Data Table 2). These divergence-driven readouts were also matched by the unadapted backbone (Supplementary Data Table 2), indicating that sensitivity to overall sequence divergence is a property of the frozen encoder, while adapter training redistributes which labels the neighbourhood resolves. Each head is therefore used for the class of task its objective targets.

### Sub-millisecond vector indexing can support CPU-tractable surveillance

To test whether MicroDUET runs on standard laboratory hardware, we profiled inference and downstream queries on a single CPU thread. The frozen backbone and both adapter heads occupy approximately 120 megabytes on disk, run within 2 gigabytes of host memory, and embed sequences at 0.28 (phylogeny) and 0.34 (function) megabases per minute on one thread of an Intel Xeon Platinum 8488C without GPU acceleration (Supplementary Data Table 1). Once embeddings are computed and stored, recurring surveillance operations such as pairwise distance estimation, nearest-neighbour retrieval and incremental database updating reduce to vector linear algebra. Across these operations, stored-embedding queries ran 10^3^-10^4^ fold faster in wall-clock time than realignment with MAFFT followed by SNP-distance calculation^32^ (Supplementary Data Fig. 6), and appending a newly sequenced isolate required sub-millisecond vector insertion rather than realignment or tree recalculation, with clade classification accuracy equivalent between the two arms. These operational characteristics demonstrate that MicroDUET can serve as an agile pre-filtering layer that triages routine sequence queries on standard laboratory hardware.

## Discussion

In this study, we present MicroDUET, a compact genomic model that supports complementary evolutionary and functional analyses of pathogen sequences. From a single frozen backbone, its two adapter heads answer the two questions surveillance asks of a sequence: the phylogeny head resolves how an isolate relates to others in circulation, and the function head resolves what it carries; gene function and resistance class. These findings show that a shared pretrained backbone can support representations tailored to distinct surveillance questions. Evaluation in *M. tuberculosis* and mpox virus further demonstrated transfer to lineage, clade, resistance and mutational-pattern tasks whose labels were absent from adapter training.

MicroDUET builds on species-aware and function-oriented genomic embedding approaches^6,13^ and complements models developed for metagenomic monitoring and viral evolutionary analysis.^33–35^ A complementary route explored elsewhere is to fine-tune a pretrained backbone on labelled data, which recovers strong taxonomic classification but updates all backbone weights and yields a separate model per task. Freezing the backbone and training small adapter heads instead keeps one shared encoder and a single embedding pass per sequence which may be more computationally efficient than a multi-model approach. Fine-tuning a backbone that also carries task-specific heads would combine the two approaches.^36^ The results for detection of rifampicin resistance illustrate why the organisation of these representations matters. In *M. tuberculosis*, resistance-associated variation can be confounded by population structure.^15^ The function head discriminated resistance without lineage-mean adjustment, whereas the phylogeny head showed substantially poorer resistance discrimination. Because the backbone was shared and frozen, this divergence demonstrates how adaptation can change which sequence relationships dominate neighbourhood-based inference. The function head nevertheless retained lineage information, indicating specialisation with overlapping biological signals. These results support reduced lineage confounding in the evaluated cohort and show that strong taxonomic retrieval alone does not establish suitability for phenotype prediction.

MicroDUET’s sensitivity to substitutions and gene loss provides a basis for studying several routes of pathogen evolution. The MPXV deletion results support screening for changes in gene content associated with genome reduction. Future work extending this approach to recombination would test whether embeddings capture mixed ancestry, while comparison with immune-recognition measurements would establish their relevance to antigenic change.

MicroDUET’s capabilities have several applications to public health surveillance. First, rapid screening against historical embeddings enables the identification of unusual sequences and emerging variants for further investigation, as demonstrated by the retrospective detection of the MPXV G.1 lineage.^37^ Second, tracking evolutionary relationships and mutational patterns, including APOBEC3-associated editing signatures, provides insights into pathogen evolution and transmission dynamics. Third, resistance-associated embeddings can help prioritise isolates for confirmatory molecular analysis or phenotypic drug susceptibility testing. Finally, MicroDUET’s computational efficiency could support timely genomic intelligence in settings with limited computing infrastructure. Once sequences have been encoded and stored, distance estimation, neighbour retrieval and database updates become vector operations, enabling queries to be performed 10^3^-10^4^ fold faster than the benchmarked realignment workflows on a single CPU thread. This comparison excludes initial sequence encoding, whose cost must be considered when processing new samples or establishing a reference database.

Several limitations define the scope of these findings. Evaluation was retrospective and focused on two pathogens, with resistance assessed from one gene per drug and discrimination varying between drugs. Neighbour-based inference depends on reference-gallery coverage, and prospective studies are needed to establish alert specificity and performance across additional pathogens and resistance mechanisms. MicroDUET requires assembled or consensus sequences and inherits their quality and preprocessing limitations. Variant-associated displacement may also reflect tokenisation effects alongside biological sequence context; its relationship to antigenic effects remains to be established, and recombination detection was not evaluated. The approach complements sequencing quality control, detailed variant and phylogenetic analyses, and confirmatory susceptibility testing. Prospective evaluation is needed to determine whether rapid embedding-based triage improves the prioritisation of genomic investigations in routine surveillance.^38,39^

In conclusion, MicroDUET demonstrates that a compact, dual-head CPU-accessible foundation model can support accurate genomic screening and rapid sequence comparisons across distinct pathogen types. By reducing computational barriers and enabling rapid genomic triage, this approach can assist with identifying threats of high public health importance in resource limited settings, providing a scalable analytical layer to strengthen global pathogen surveillance.

## Competing Interests Policy

I declare that the authors have no competing interests or other interests that might be perceived to influence the results and/or discussion reported in this paper.

## Data Availability Statement

Data available on request

## Acknowledgements

We gratefully acknowledge the assistance of Sequentia Biotec, in particular Daniel Julián Alonso and Luis García Jimeno, for their assistance with bioinformatic processing of the *M. tuberculosis* and MPXV datasets. This study was partially supported by the University of Sydney – National University of Singapore Ignition Grants.

The authors would like to thank the Gates Foundation (INV-037608, INV-064510) for supporting the establishment of the Asia Pathogen Genomics Initiative. We also appreciate the support of the Gates Foundation (INV-079147), Temasek Foundation, the Philanthropy Asia Alliance, and the University of Sydney – National University of Singapore Ignition Grants for our work on artificial intelligence and pathogen genomics.

## Methods

### Two-head embedding architecture

The two adapter heads share a single frozen ProkBERT-mini-long backbone encoder^11^ (six-mer tokenization with a stride of two, hidden dimension 384, 26.6 × 10⁶ parameters), whose weights remained fixed throughout all training stages. Each task-specific head comprises a two-layer pre-layer-normalization Transformer encoder (six attention heads, feed-forward intermediate dimension 1,536, Gaussian error linear unit activations, and dropout probability 0.1; 3.5 × 10⁶ parameters each) operating directly on the backbone encoder’s final-layer token representations. Token states are aggregated via masked mean pooling across non-padding positions and L2-normalized to yield a 384-dimensional unit vector, z. Neither head outputs a class label. Labels (species, gene function, resistance class, lineage, clade) are assigned by cosine nearest-neighbour retrieval against a labelled reference set embedded by the same head (k = 1 for the retrieval benchmarks; k = 5 for lineage, clade and resistance classification). For sequences exceeding the 4,096-bp context window, sequences are partitioned into 4,096-bp windows with a 60-bp overlap; window embeddings are averaged and re-normalized to unit Euclidean length. A single forward pass through the frozen backbone simultaneously generates representations for both adapter heads. Both heads were optimized on a single NVIDIA L4 GPU using AdamW^40^ (learning rate 1.73 × 10⁻⁴, weight decay 0.01), batch size 48, bfloat16 mixed precision, and gradient norm clipping at 1.0. The learning rate warmed up linearly over the initial 2.5% of optimization steps before decaying under a cosine schedule. Validation loss was evaluated every 500 steps; the checkpoint achieving optimal validation performance was retained, with training terminated after 20 consecutive evaluations without validation metric improvement greater than 10⁻³.

### Species-identification adapter head training

The species-identification phylogeny head was trained to optimize taxonomic sequence-origin discrimination using a symmetric InfoNCE contrastive loss^41^ at temperature τ = 0.05 applied to a training-only projection head (384 → 384 [ReLU] → 128, L2-normalized), which was discarded after training. Positive pairs comprised two non-overlapping windows extracted from the same genome; the remaining representations within the batch served as negative comparators. Cellular pairs were drawn from the 2,000,000 DNABERT-S training pairs,^13^ consisting of approximately 10-kb genomic segments from GenBank bacterial and fungal genomes. Viral pairs were sampled from complete NCBI RefSeq viral genomes.^42^ To prevent evaluation leakage, genomes matching the 1,357 evaluation taxonomy identifiers were strictly excluded prior to training. 5% of pairs were held-out for validation and evaluated by InfoNCE loss.

### Gene-function adapter head training

The gene-function function head was trained to capture functional orthology across divergent evolutionary lineages using a normalized-softmax cross-entropy classification loss at temperature τ = 0.05. Embeddings were multiplied by an L2-normalized weight matrix for each respective training collection (logits = cosine similarity / 0.05); cross-entropy was computed over that dataset’s classes and averaged across the batch, with classifier weights discarded after training. Bacterial and archaeal coding sequences were obtained from the Scorpio gene–taxa collection.^6^ 5% of sequences were held-out as a validation set. We sampled classes proportional to the square root of their record count; taxon uniformly within the class; and sequence uniformly within the taxon. Batches were drawn from 16-batch buffers sorted by sequence length to minimize padding. Optimization was conducted using AdamW with identical optimizer hyperparameters, mixed precision, and learning rate schedules as the phylogeny head, maintaining a frozen backbone encoder throughout.

### Viral orthologue corpus and validation schedule

Viral coding sequences were obtained from VOGDB release 237.^43^ Poxviridae were strictly excluded to prevent task leakage. Redundancy was collapsed via MMseqs2 linclust^44^ at 95% nucleotide identity and 80% coverage, retaining one sequence per cluster within each orthologous group. Groups containing at least 10 coding sequences across at least five viral taxa were retained. Groups were partitioned into 10% test, 5% validation, and 85% training splits. For validation, classes were capped at 100 (Scorpio^6^) or 20 (VOGDB^43^) records per class, yielding 2,200 records in 22 classes and 4,181 records in 266 classes with zero overlap with training classes. Validation embeddings were evaluated every 500 optimization steps by leave-one-out cosine k-nearest-neighbour classification (k = 5, inverse cosine distance weighting).

### Species-identification benchmark

To measure whole-genome sequence-origin discrimination, models were evaluated on 1,920 held-out sequence windows (1,000 cellular windows across 200 species; 920 viral windows across 200 species) spanning lengths from 100 bp to 10 kb under the evaluation framework established by Zhou et al.^13^ Windows were sampled from GenBank bacterial and fungal genomes for cellular taxa, and from NCBI RefSeq viral genomes^42^ for viral taxa. As detailed above, all candidate genomes matching the 1,357 evaluation taxonomy identifiers were strictly excluded prior to model training to exclude these taxa from adapter training. Performance was evaluated via leave-one-out one-nearest-neighbour cosine retrieval of the matching species label, where a successful retrieval required the top-ranked neighbour to share the query organism’s species designation (the species name recorded for the reference sequence; no other taxonomic rank was scored).

### Gene-function benchmark

To measure recovery of functional labels on held-out classes from the Function-head training distribution, models were evaluated on 2,200 coding sequences (1,200 cellular sequences across 60 Scorpio gene labels; 1,000 viral sequences across 100 VOGDB orthologous groups). Cellular sequences were drawn from the Scorpio gene-out set, whose 60 labels are absent from the Scorpio training data, with 20 sequences per label and one sequence per genus (189–3,396 bp; median 988 bp)^6^. Viral sequences were drawn from the 10% VOGDB release 237 group split withheld from training and from checkpoint selection^43^, after the 95% identity collapse described above, with 10 sequences per group and one sequence per viral taxonomy identifier (150–6,963 bp; median 444 bp). Coding sequences shorter than 150 bp, longer than 12,288 bp, or containing ambiguous nucleotides were excluded. A neighbour sharing the query genus (cellular) or taxonomy identifier (viral) was excluded. Performance was evaluated via one-nearest-neighbour cosine retrieval on L2-normalized embeddings (k = 1), where a successful retrieval required the top-ranked permitted neighbour to share the query’s gene label (cellular) or orthologous-group label (viral).

### Resistance-class retrieval benchmark

To evaluate whether sequence representations place functionally related coding sequences together outside the training distribution, we assembled a benchmark independently of the training data, consisting of NCBI AMRFinderPlus reference gene catalogue.^20^ Each coding sequence was labelled with the functional resistance or stress class from its reference protein record and assigned to a gene family by its parent node in the AMRFinderPlus hierarchy. To exclude sequences encountered during training, both gene-function training corpora were searched against all 8,881 catalogue sequences using MMseqs2 nucleotide search^44^ (sensitivity 7.5, at most 50 target sequences per query, E ≤ 10⁻⁵). Any catalogue sequence matching at ≥ 80% identity over ≥ 80% of its length was excluded (287 sequences removed). Classes with at least 10 remaining gene families were retained, and up to 20 coding sequences per class were sampled, one per family. The final panel contains 398 coding sequences from 23 classes and 398 families (lengths 201–3,231 bp, median 798 bp). Performance was evaluated via one-nearest-neighbour cosine retrieval on L2-normalized embeddings (k = 1), where a successful retrieval required the top-ranked neighbour to share the query’s resistance or stress class.

### Benchmark baselines and reference alignment

Representation quality across screening benchmarks was evaluated against analytical chance (1.67% cellular and 1.00% viral on gene function; 4.35% on resistance class), an oligonucleotide six-mer composition cosine baseline under the same k = 1 nearest-neighbour rule, and MMseqs2 translated local alignment search^44^ (sensitivity 7.5, at most 2,000 target sequences per query, E ≤ 10), taking the best-bitscore permitted target. Translated alignment defines the empirical ceiling for sequence homology search rather than a competing embedding method.^44^ Oligonucleotide frequency vectors were constructed by counting canonical six-mer occurrences across sequences and L2-normalizing to unit Euclidean length prior to cosine distance retrieval. Each model was compared with native ProkBERT-mini-long^11^ by a paired exact McNemar test^45^ on per-query hits, applying Holm step-down multiplicity correction^46^ across models and both reference methods within each view (cellular gene function, viral gene function, and resistance class).

### Clinical Mycobacterium tuberculosis cohort bioinformatic analysis

We selected 1,766 clinical *M. tuberculosis* isolates from the CRyPTIC consortium compendium^28,29^ which had whole genome sequences and associated phenotypic drug susceptibility test (DST) results using the CRyPTIC consortium standardised panel. Raw paired-end Illumina sequencing reads were obtained from the provided Sequence Read Archive accession numbers. We performed variant calling using the TBProfiler 6.7.0 pipeline,^47^ retaining variants with at least 50% frequency and 10x depth after requiring a minimum post-trimming read length of 30 and Phred score of 20. Using samtools 1.20 and bcftools 1.20, we excluded samples where the mean read depth was below 20x and generated a consensus genome for each sequence by applying the filtered variants to the reference sequence H37Rv (RefSeq GCF_000195955.2; GenBank AL123456.3).^48^ We then masked any position with a depth below 10x or an ambiguous nucleotide character. Genes that were at least 90% masked were deleted from the consensus sequence provided that they were genes where this occurred for at least 5% of sequences. We then excluded variants in masked positions. We retained the whole consensus sequence, consensus sequences for a selection of 39 drug resistance associated genes, and TBProfiler’s genomic DST call for rifampicin.

### Mpox virus cohort bioinformatic analysis

We obtained 11,266 assembled MPXV genomes from the Nextstrain mpox builds, restricted to sequences deposited without embargo. We used the nextclade mpox clade-specific workflows to produce a quality-controlled multiple sequence alignment.^27,38^ We applied the masks included in the nextclade workflow and additionally masked all positions with ambiguous nucleotide characters. We removed gaps and applied indels where they occurred outside of masked regions. Genes that were at least 90% masked were deleted from the consensus sequence provided that they were genes where this occurred for at least 5% of sequences. Spans that met this criterion but were also part of a retained gene were themselves retained. We counted the number of substitutions with respect to the reference genomes (KJ642613.1 for clade I and NC_063383.1 for clade II), aggregating by nucleotide context to distinguish substitutions consistent with APOBEC-3 edits (i.e. TC→TT and GA→AA).^30^ Finally, we imputed the value of all masked positions using the value at the corresponding position in the clade-specific reference genome. We retained whole genome consensus sequences, gene-level consensus sequences, and predicted lineage and clade from the nextclade pipeline.

### In silico mutational trajectory simulations

To evaluate whether continuous cosine distance preserves discrete mutational separation, we simulated independent stochastic mutational trajectories across two benchmark loci: *M. tuberculosis rpoB* (3,519 bp, H37Rv reference strain,^48^ GenBank AL123456.3) and mpox virus OPG210 (5,643 bp coding sequence from isolate PP_000SW2F; GenBank PQ240044.1). For each gene, 30 independent stochastic trajectories were simulated across 14 discrete single-nucleotide polymorphism (SNP) substitution counts (k ∈ {1, 2, 5, 10, 15, 20, 25, 30, 40, 50, 64, 78, 100, 150} substitutions; n = 420 pairs per gene). Single-nucleotide substitutions were introduced stochastically without replacement across mutable positions. Pairwise cosine embedding distances (defined as 1 − u · v for L2-normalized vectors u and v) were computed relative to the unmutated reference sequence. Association between embedding distance and mutational distance was quantified using Spearman rank correlation coefficients (ρ) with replicate-bootstrap 95% confidence intervals and two-sided significance tests with Holm correction^46^ across substitution counts.

### In silico coding variant consequence profiling

To test whether sequence representations differentiate annotated variant classes, we generated in silico coding variants across the 4,096-bp context windows of *M. tuberculosis rpoB* (*n* = 4,626 variants) and mpox virus OPG210 (isolate PP_000SW2F; GenBank PQ240044.1; *n* = 2,988 variants). Every GC to AT substitution in the window was introduced, together with a draw of 50 frameshift insertions (1 bp or 2 bp) and 50 in-frame insertions or deletions per gene. Variants were classified with the standard genetic code into synonymous, non-synonymous, and nonsense substitutions; the initiation codon was held out, and frameshift insertions were split into +1 and +2. L2-normalised cosine distance from the unmutated sequence was compared among the six classes by a Kruskal–Wallis test and by two-sided Dunn tests on that pooled ranking, Holm-adjusted across the 15 pairs at *α* = 0.05. A shared letter means the classes were not statistically separable.^49^

### Clinical *M. tuberculosis* population structure

To evaluate population structure resolution in clinical isolates, we examined the 3,638 unique coding sequences across the gene-level consensus sequence dataset. Global sequence distribution across loci was visualized using two-dimensional UMAP^24^ (15 neighbours, minimum distance 0.1, cosine metric). Lineage structure was evaluated on *rpoB* coding sequences across 347 clinical isolates with unambiguous lineage annotations^50^ spanning major lineages 2 (East Asian/Beijing, n = 79), 3 (Central Asian, n = 137), and 4 (Euro-American, n = 131). Neighbourhood cohesion was quantified via leave-one-out nearest-neighbour analysis (k = 5, inverse cosine distance weighting). Performance at that choice was stable for k = 1-20 (Supplementary Data Fig. 7a-c).

### Rifampicin resistance prediction and lineage deconvolution

Rifampicin resistance was evaluated across 347 *rpoB* consensus sequences from the gene level dataset. Out-of-fold five-fold cross-validated nearest-neighbour classification was performed at *k* = 5 using inverse cosine distance weighting across training folds. Naive resistance scores were evaluated by area under the receiver operating characteristic curve (AUROC) and misclassification counts. To decouple resistance from phylogenetic background, the lineage-mean vector was subtracted from embeddings and rifampicin resistance via *k*-nearest neighbours recalculated.

### Multi-drug resistance discrimination

Consensus sequences of one gene most associated with resistance for each drug (*rpoB*, *katG*, *pncA*, *embB*, *gyrA*, Rv0678, *ethA*, *gid*) were embedded with the function head and scored as for rifampicin (out-of-fold five-fold cross-validated nearest-neighbour classification, k = 5, inverse cosine distance weighting). Neighbour weight is the share of that weighting carried by resistant neighbours, and discrimination was summarised by AUROC. Resistance was defined either from TB-Profiler calls alone or from CRyPTIC phenotypic DST or a TB-Profiler call. Isolates with resistance called at a different gene were retained as resistant.

### Clinical mpox virus cohort and quarterly lineage dynamics

Intra-clade epidemic dynamics were evaluated on 8,794 whole-genome consensus sequences from mpox virus clade IIb from curated Nextstrain builds.^27^ Quarterly lineage mix was scored on the 8,738 of these collected in the 16 quarters from 2022-Q2 to 2026-Q1. Whole-genome embeddings were generated using the phylogeny head across tiled 4,096-bp windows with 60-bp overlap, followed by mean-pooling and L2-normalization. Out-of-fold nearest-neighbour classification (*k* = 5) was performed to predict circulating sublineages (B, C, F, and G). Concordance between observed and embedding-predicted quarterly lineage proportions was the total variation distance between the two mixes, averaged across quarters with weights proportional to the number of isolates in each quarter. The 95% confidence interval is a 2,000-draw bootstrap. A quarter was counted as within sampling error when that distance fell below the 95th percentile of distances from resampling that quarter’s isolates alone.

### Emergent viral lineage anomaly detection

To evaluate unindexed variant detection, 266 whole-genome consensus sequences of the emergent 2025 mpox virus lineage G.1 from Sierra Leone^18^ were evaluated against a historical gallery of 3,842 clade IIb whole genomes from the 2022 B.1 epidemic, accompanied by 266 held-out 2022 B.1 genomes as in-distribution controls. Anomaly scores were defined as the cosine distance to the fifth nearest neighbour in the static gallery. Discrimination of G.1 from in-distribution controls was evaluated by AUROC with two-sided DeLong tests^51^ against analytical chance (0.5). Quarterly emergence dynamics were monitored by scoring post-2022 circulating clade IIb genomes (non-G.1 clade IIb genomes, n = 1,392) from 2023-Q1 to 2024-Q4 alongside G.1 arrivals across 2025-Q1 (n = 19), 2025-Q2 (n = 193), and 2025-Q3 (n = 53).

### APOBEC3 mutational signature analysis and evolutionary era reconstruction

To evaluate sensitivity to host-mediated editing, 30 independent stochastic trajectories were simulated up to 150 substitutions, comparing canonical APOBEC3 transitions (TC→TT and GA→AA)^22,30^ against equal counts of GC-matched non-motif substitutions. Differences in mean cosine displacement were evaluated across 14 substitution levels using two-sided Mann–Whitney U tests with Holm correction.^46^ Across 8,563 clinical mpox virus consensus genomes, observed genomic proportions of APOBEC3 mutations were quantified from variant call records relative to the reference genome [KJ642613.1 for clade I and NC_063383.1 for clade II]. The APOBEC3-edit proportion was predicted by leave-one-out cosine nearest-neighbour regression (k = 5, inverse-distance weights). R² is the coefficient of determination between the observed and predicted proportions; the p-value is their Pearson correlation.

### Operational hardware benchmarking

Computational efficiency was profiled on a single physical thread of an Intel Xeon Platinum 8488C CPU (2.40 GHz, 32 GB RAM). Three routine surveillance operations were benchmarked across cohort sizes N ∈ {16, 32, 64, 128, 256, 512, 1024, 2048, 4096, 8192, 11129} (capped at 2,048 for alignment): pairwise distance matrix estimation, nearest-neighbour retrieval (k = 10) and incremental database insertion.^52^ The alignment arm utilized MAFFT (--thread 1 --retree 2 --maxiterate 0) for multiple sequence alignment, then a SNP-distance matrix. The embedding arm utilized BLAS matrix multiplication for distance calculation, PyNNDescent (version 0.5.11)^52^ for nearest-neighbour indexing and vector appending for incremental updates. Execution times were recorded as median wall-clock duration across five iterations following warm-up, and memory was measured as peak resident set size.

### Statistical reporting and reproducibility

All statistical tests were two-sided unless explicitly stated otherwise. Significance thresholds were set at α = 0.05, with multiple testing corrections applied via the Holm step-down procedure^46^ across all multi-comparison screens. Non-parametric rank tests (Spearman rank correlation ρ, Mann–Whitney U test,^53^ Kruskal–Wallis omnibus test,^49^ Dunn pairwise test^54^) were employed for mutational distances, consequence displacements, and non-normally distributed metrics. Confidence intervals for classification accuracies and retrieval rates were computed using binomial Wilson score intervals^55^ with continuity correction where appropriate. Confidence intervals for rank correlations were computed via 2,000 replicate bootstrap resamples across trajectories. DeLong tests^51^ were used for correlated and single-sample ROC curve comparisons. Regression goodness of fit was evaluated using the coefficient of determination (R²) between observed and predicted values, with significance from Pearson correlation of the two series.^56^

## Notes

### Competing Interest Statement

The authors have declared no competing interest.

