## Supplementary Data for "MicroDUET: a dual-headed genomic foundation model for pathogen surveillance"

**Supplementary Data Table 1** | Foundation model screen across species discrimination and inference throughput. Benchmark of 17 pretrained genomic foundation models and MicroDUET. Benchmarks and megabase throughput measures are outlined in methods; throughput is the mean of 3 measures. Bolded numbers represented the best score per column.

| Model | Params (N) | SpeciesID (cellular) | SpeciesID (viral) | AMRFinderPlus | FunctionID (cellular) | FunctionID (viral) | CPU Mb/min (XEON) | GPU Mb/min (L4) |
|---|---|---|---|---|---|---|---|---|
| ProkBERT[11] | **26.6M** | 57.9 | 61.7 | 56.3 | 37.4 | 83.9 | 0.409 | 13.477 |
| MicroDUET (head) | 33.6M | **80.3 (phylogeny)** | **66.6 (phylogeny)** | **74.4 (function)** | **97.3 (function)** | **97.2 (function)** | 0.281 | 21.92 |
| DNABERT-S[13] | 117M | 77 | 59.9 | 39.4 | 13.8 | 63.3 | 0.489 | 17.271 |
| NT-v2-50M[8] | 55.9M | 48.6 | 60.3 | 50 | 40.1 | 73.3 | 0.055 | 2.122 |
| GENA-BERT[57] | 135M | 48.5 | 56.5 | 45.5 | 31.2 | 72.1 | 0.6 | 22.101 |
| NT-v2-50M-3mer[8] | 50.0M | 47.5 | 59.1 | 54.5 | 36.8 | 76.9 | 0.056 | 2.083 |
| GenomeOcean-100M[35] | 120M | 43.3 | 57.4 | 43.5 | 34.9 | 61.2 | 0.635 | 20.503 |
| GenomeOcean-4B[35] | 4.25B | 42.8 | 60.1 | 46 | 33.4 | 61.6 | 0.023 | 1.284 |
| NT-v2-500M[8] | 498M | 41 | 53.4 | 42 | 32.4 | 67.7 | 0.018 | 0.684 |
| NT-2.5B[8] | 2.50B | 37 | 54.3 | 52.8 | 50.2 | 72.5 | 0.005 | 0.29 |

| | | | | | | | | |
|---|---|---|---|---|---|---|---|---|
| METAGENE-1[34] | 6.48B | 36.1 | 46 | 37.4 | 20.2 | 60.7 | 0.008 | 0.44 |
| METAGENE-1 8-bit[34] | 6.49B | 36.1 | 46.1 | 37.9 | 21.1 | 59.4 | NA | 0.44 |
| DNABERT-2[58] | 117M | 32.4 | 47.6 | 45 | 31.9 | 65.5 | 0.389 | 14.257 |
| GENA-BigBird[57] | 114M | 32.1 | 60.4 | 36.9 | 19.8 | 61.4 | 0.629 | 18.865 |
| Evo2-7B-base[7] | 7B | 24.4 | 49.8 | 43.5 | 31.3 | 59.4 | NA | 0.16 |
| Evo2-7B[7] | 7B | 24 | 50.7 | 42.5 | 32.5 | 62 | NA | 0.169 |
| PathoLM[59] | 55.9M | 17.7 | 15.3 | 17.3 | 7.1 | 17.5 | **1.257** | **28.821** |
| GENERanno-0.5B[60] | 493M | 14.1 | 53.7 | 47.5 | 49.8 | 58.8 | 0.027 | 1.272 |

**Supplementary Data Table 2**. | A comparison of Native ProkBert (ProkBERT-mini-long) vs the dual heads of MicroDUET. Bolded numbers represented the best score per row.

| **Benchmark** | **ProkBERT** | **Phylogeny** | **Function** |
|---|---|---|---|
| SpeciesID cellular, (k = 1) | 0.579 | **0.803** | 0.316 |
| SpeciesID viral, (k = 1) | 0.617 | **0.666** | 0.643 |
| Gene function ID, cellular (k=1) | 0.374 | 0.141 | **0.973** |
| Gene function ID, viral (k=1) | 0.839 | 0.695 | **0.972** |
| AMRFinderPlus (k=1) | 0.563 | 0.452 | **0.744** |
| OPG210 embedding–SNP Spearman ρ (n = 420) | 0.944 | **0.964** | 0.948 |
| MTB lineage neighbour weight (n = 347) | 0.917 | **0.943** | 0.604 |
| Lineage 2 neighbour weight (n = 79) | 0.836 | **0.890** | 0.374 |
| OPG210 clade neighbour weight (n = 404) | 0.973 | **0.982** | 0.978 |
| MPXV Clade balanced accuracy (n = 404) | 0.924 | 0.975 | **0.987** |
| Clade IIa recall (n = 5) | 0.8 | **1** | 1 |
| MPXV Lineage TVD error (pp) | **1.2** | 1.3 | 1.5 |
| MTB Lineage balanced accuracy (n = 347) | **0.966** | 0.965 | 0.684 |
| Rifampicin naive AUROC (n = 347) | 0.876 | 0.735 | **0.985** |
| Rifampicin k = 5 errors / 347 | 41 (33 FP, 8 FN) | 52 (45 FP, 7 FN) | **22 (4 FP, 18 FN)** |
| Whole-genome embedding: 2025 G.1 vs 2022 B.1 gallery AUROC | 0.978 | **0.981** | 0.978 |
| MPXV G.1 AUROC, 2025-Q1 (n = 19) | 0.967 | **0.974** | 0.966 |

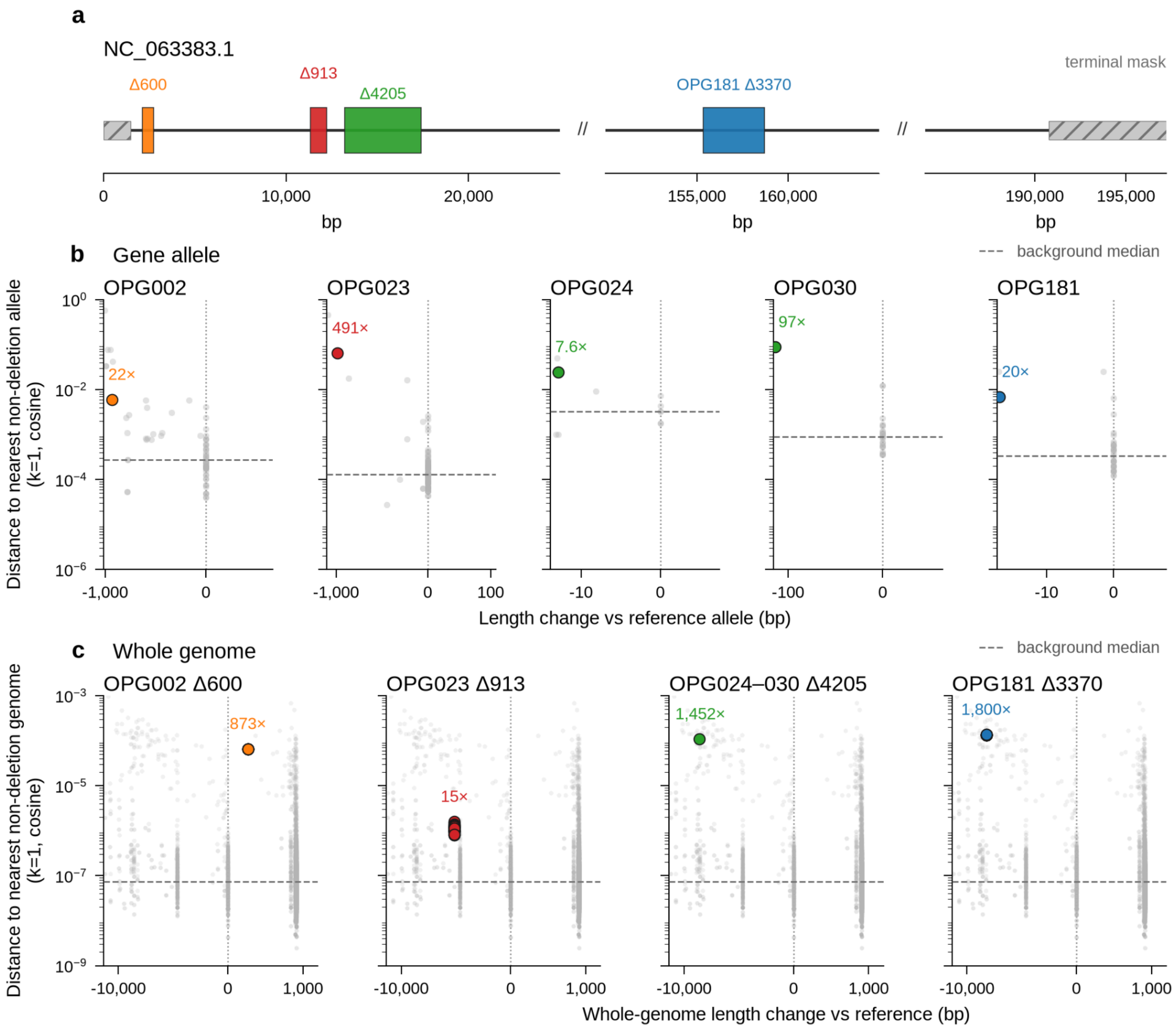


**Supplementary Data Fig. 1** | Deletion-associated embedding displacement at gene and whole-genome scales in mpox virus. **a**, Schematic of four deletion events in the MPXV reference genome NC_063383.1: Δ600, Δ913, Δ4205 (spanning OPG024–OPG030) and OPG181 Δ3370 . Hatched regions indicate terminally masked sequence; double slashes indicate omitted genomic intervals. **b,** Cosine distance to the nearest non-deletion allele (k = 1) for five affected gene alleles (OPG002, OPG023, OPG024, OPG030 and OPG181), plotted against allele-length change relative to the reference. **c**, Cosine distance to the nearest non-deletion whole genome embedding (k = 1) for genomes carrying the indicated deletions, plotted against whole-genome length change relative to the reference. Grey points show background non-deletion sequences, and dashed horizontal lines indicate the background median distance. Coloured points highlight deletion variants; labels indicate their distance relative to the background median.

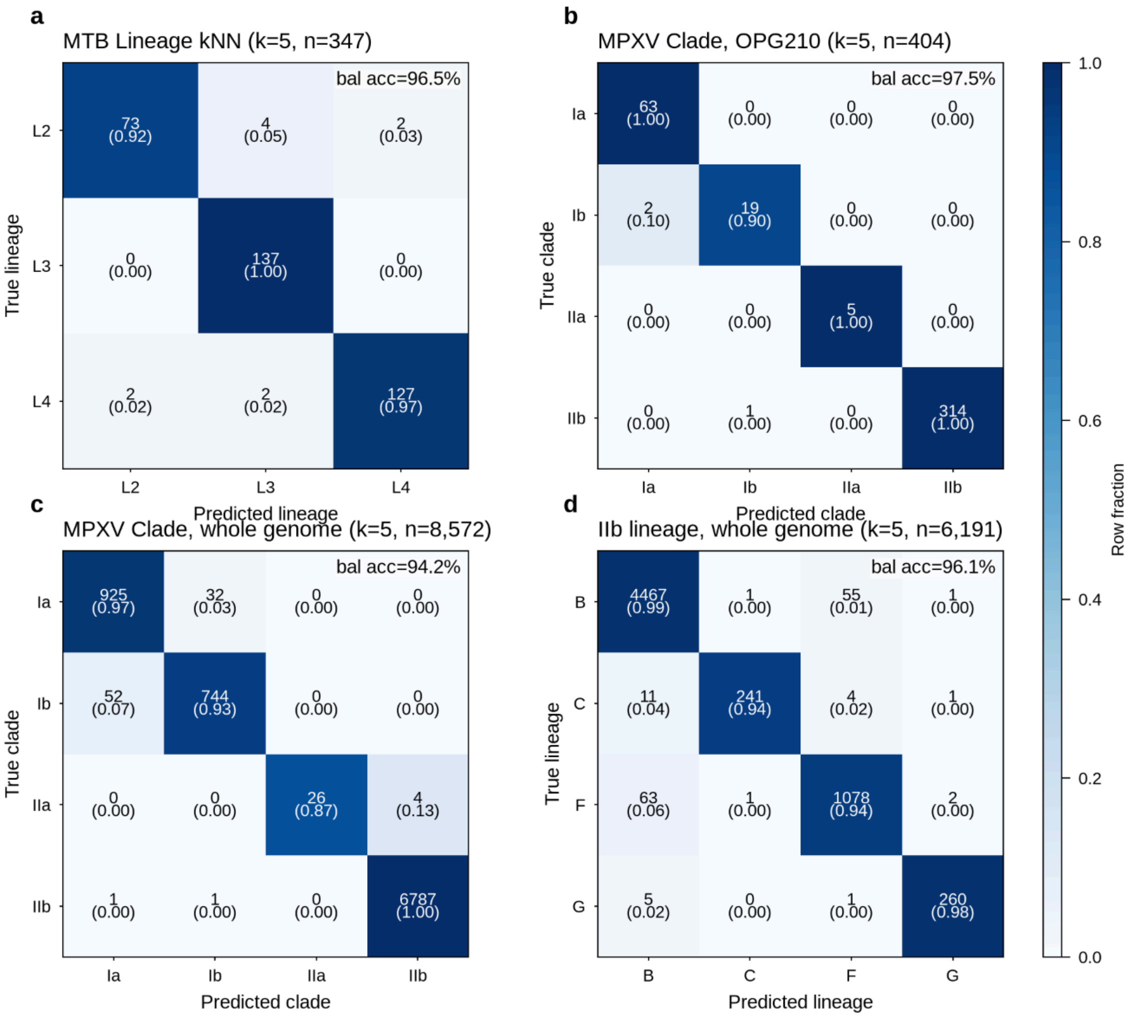


**Supplementary Data Fig. 2** | Confusion matrices for the classifications summarised in Fig. 3. Leave-one-out kNN (k = 5), rows normalised to the true class; balanced accuracy is shown in each panel. **a**, MTB lineage (Fig. 3b). **b**, MPXV clade from OPG210 (Fig. 3c). **c,** MPXV clade from whole genomes. **d,** IIb lineage from whole genomes, the per-genome assignments behind the mix in Fig. 3d–e.

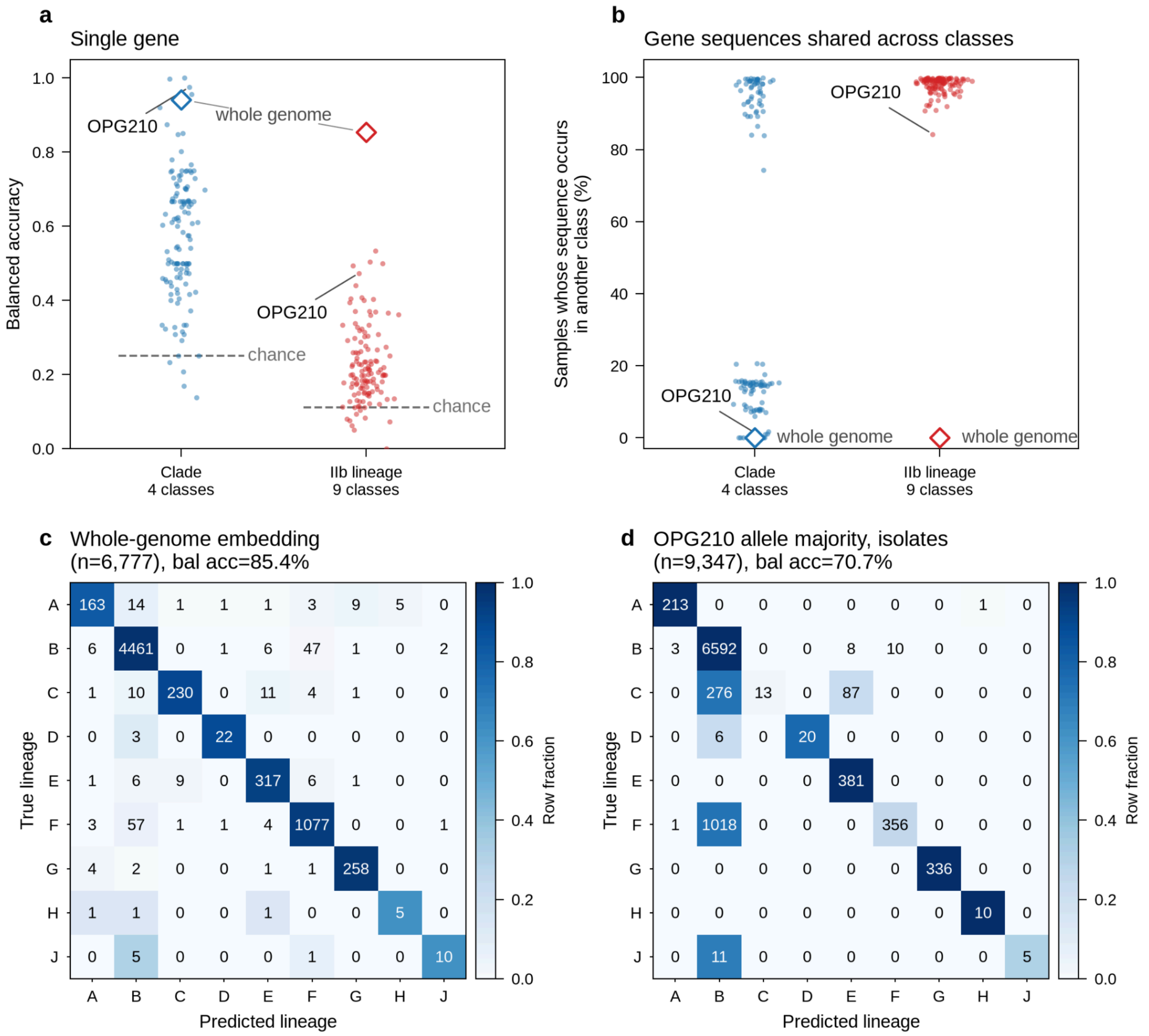


**Supplementary Data Fig. 3** | A single mpox gene resolves clade but not IIb lineage. Leave-one-out kNN (k = 5) on MicroDUET embeddings, scoring each of 124 genes and the whole genome. **a,** Balanced accuracy per gene (points) and whole genome (diamond). For clade, OPG210 (0.975) matches the whole genome (0.942); for nine IIb lineages, no gene approaches the whole genome (0.854; median gene 0.20, OPG210 0.47). **b,** Samples whose exact gene sequence occurs in another class: median 16% for clade, 99% for lineage (Wilcoxon $P = 1.1 \times 10^{-21}$). **c,** Whole-genome lineage confusion matrix behind a; errors mostly place small lineages into B. **d,** Model-free baseline with each isolate's lineage predicted based on the majority lineage of its exact OPG210 allele (0.708); lineages C and F are mostly indistinguishable using OPG210 alone.

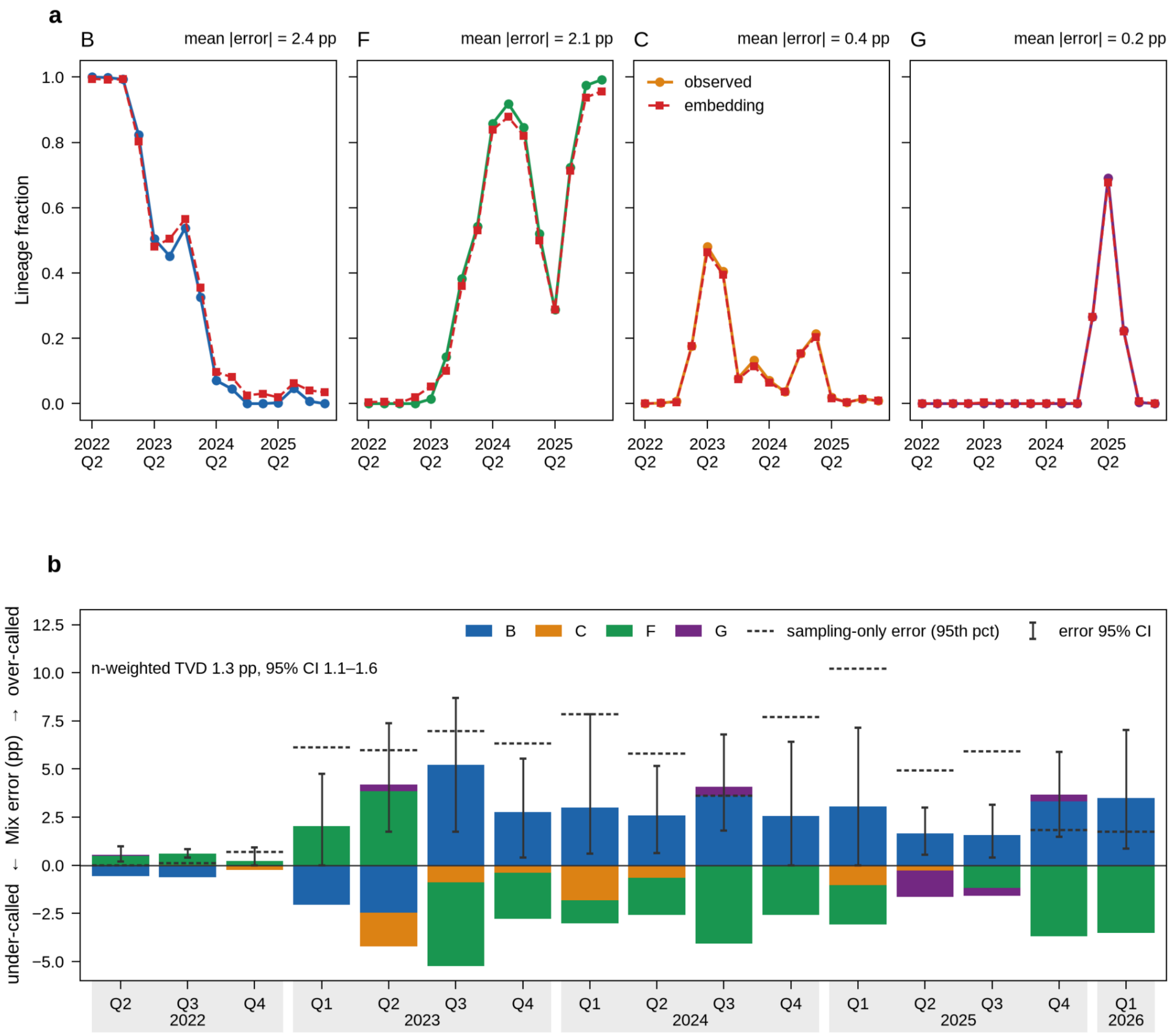


**Supplementary Data Fig. 4** | Whole-genome embeddings reconstruct the quarterly clade IIb lineage mix. Lineages (B, C, F, G) of 8,738 IIb genomes were assigned by leave-one-out kNN (k = 5) on embeddings and summed per quarter. **a**, Observed (circles) and embedding-derived (squares) lineage fractions, with mean absolute error per lineage. **b**, Signed per-quarter error (above zero = over-called) with 95% bootstrap CIs. Dashed ticks show the error expected from sampling that quarter's isolates alone. Mean error is 1.3 pp (95% CI 1.1–1.6), within sampling error in 11/16 quarters.

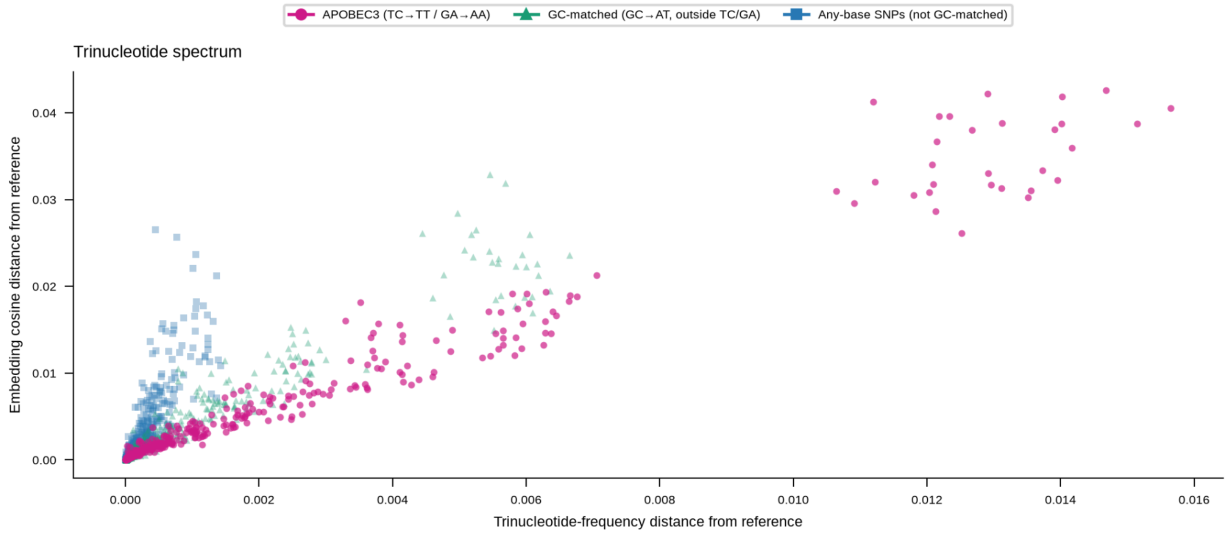


**Supplementary Data Fig. 5 |** Embedding distance tracks 3-mer frequency shift, with APOBEC3 substitutions generating the largest displacement in embedding distance. Simulated OPG210 variants plotted as embedding cosine distance from the reference against the cosine distance between 3-mer frequency vectors (all 64 overlapping trinucleotides, expressed as proportions). APOBEC3-style TC→TT / GA→AA substitutions reach a mean 3-mer distance of 0.0129, versus 0.0056 for GC-matched GC→AT controls.

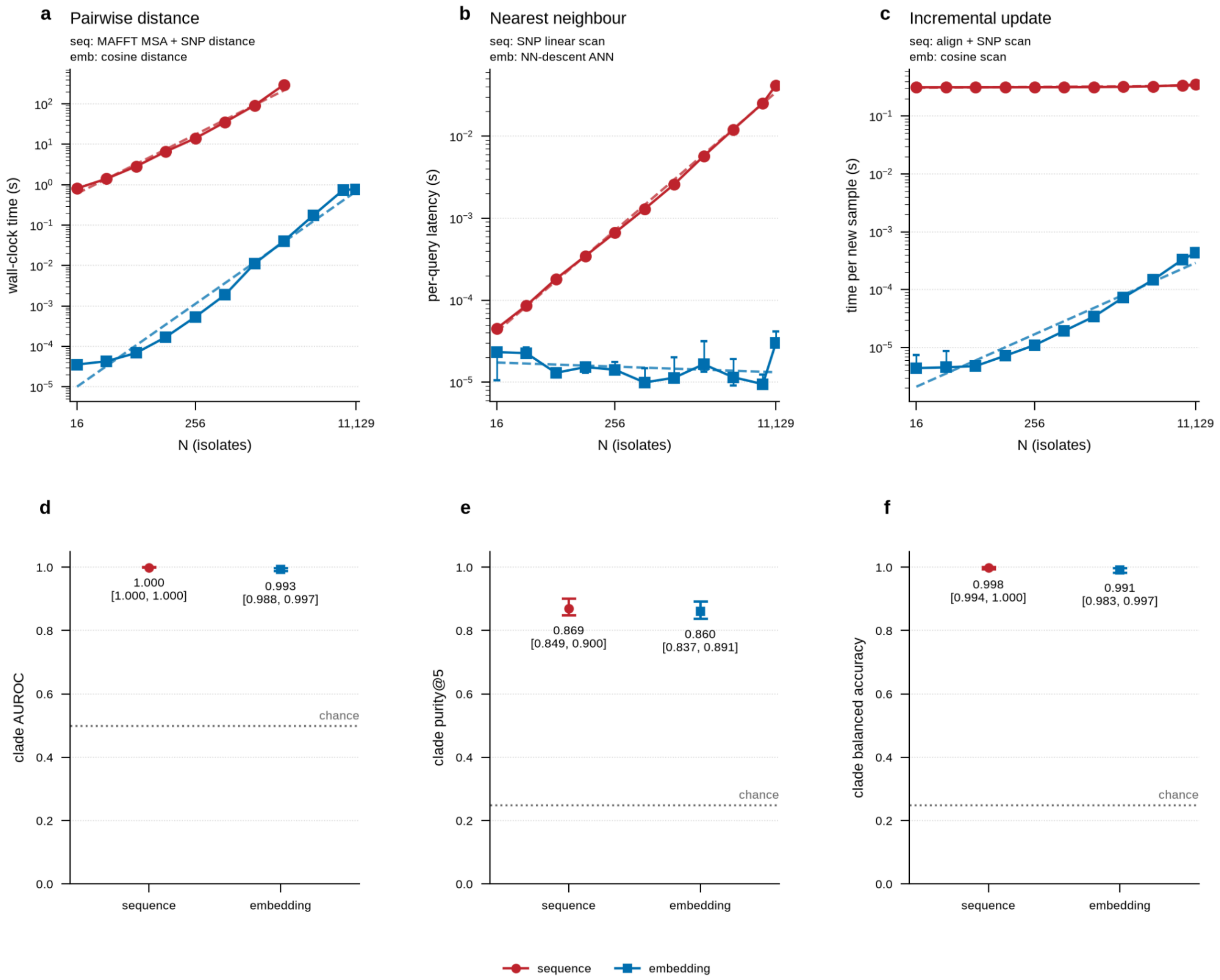


**Supplementary Data Fig. 6 | a-c**. Operational benchmarking of stored vector embeddings versus sequence-alignment workflows. Wall-clock execution time across pairwise distance estimation, nearest-neighbour retrieval, and incremental database updating on standard CPU hardware. **d–f,** Clade AUROC, nearest-neighbour purity (k = 5), and balanced accuracy for alignment versus stored embeddings.

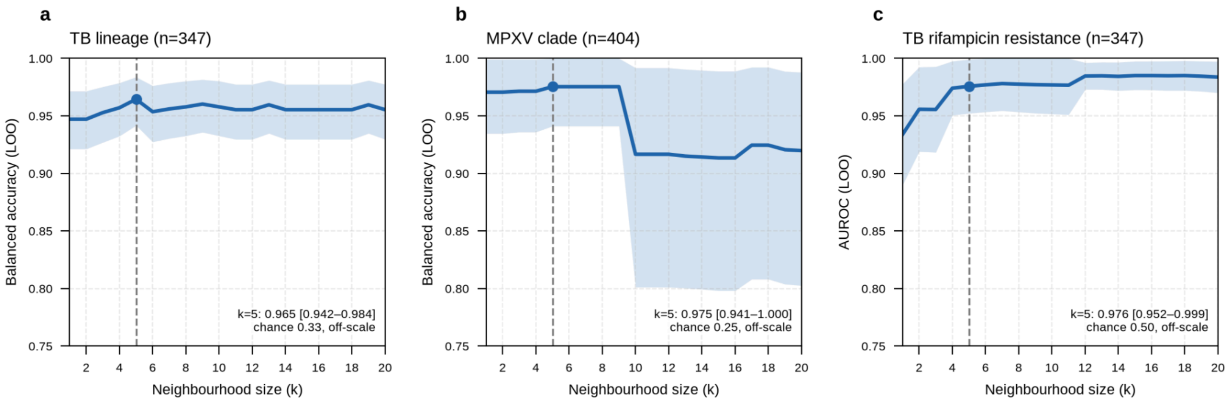


**Supplementary Data Fig. 7** | Neighbourhood results are robust to the choice of k. Leave-one-out kNN performance for k = 1–20; shaded bands are 95% CIs (bootstrap for balanced accuracy, DeLong for AUROC); dashed line marks k = 5, used throughout. **a**, MTB lineage balanced accuracy (0.947–0.965). **b**, MPXV clade balanced accuracy at OPG210. **c,** Rifampicin resistant-neighbour AUROC (≥0.95 for k ≥ 2).

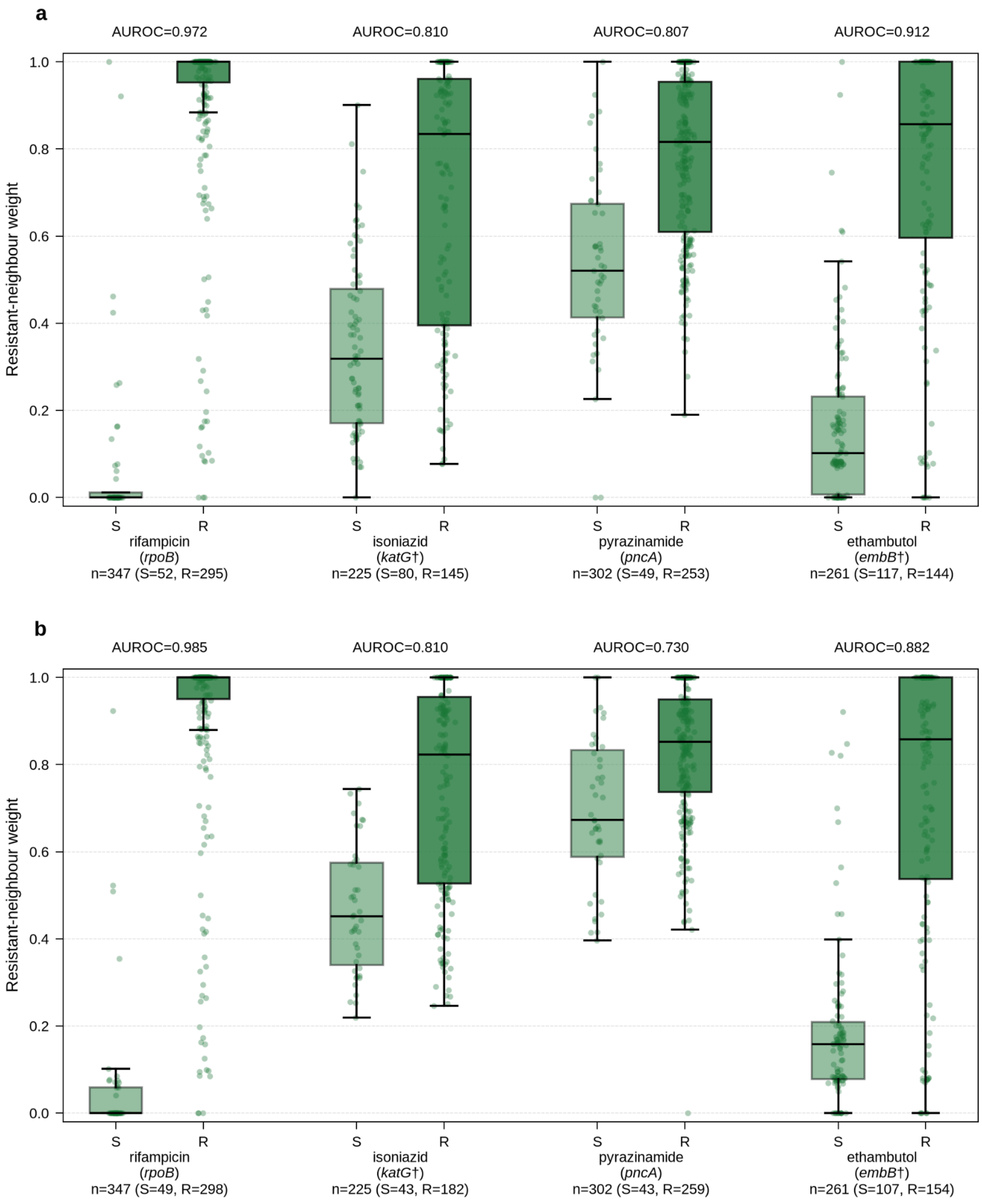


**Supplementary Data Fig. 8** | MicroDUET function-head neighbourhoods discriminate resistance for first-line antitubercular drugs. For each drug, the gene shown was embedded with the function head, and each isolate was scored by the neighbour weight on resistant isolates (out-of-fold, k = 5, inverse cosine distance weighting; y-axis). Points are isolates; boxes show the median and interquartile range, whiskers extend to 1.5× the interquartile range. S, susceptible; R, resistant; AUROC is shown above each pair. **a,** TB-Profiler resistance calls only. **b,** CRyPTIC phenotypic DST or a TB-Profiler resistance call. †, some resistant isolates are called at a different gene from the one plotted (panel b): isoniazid 27 of 182, ethambutol 4 of 154.**f**

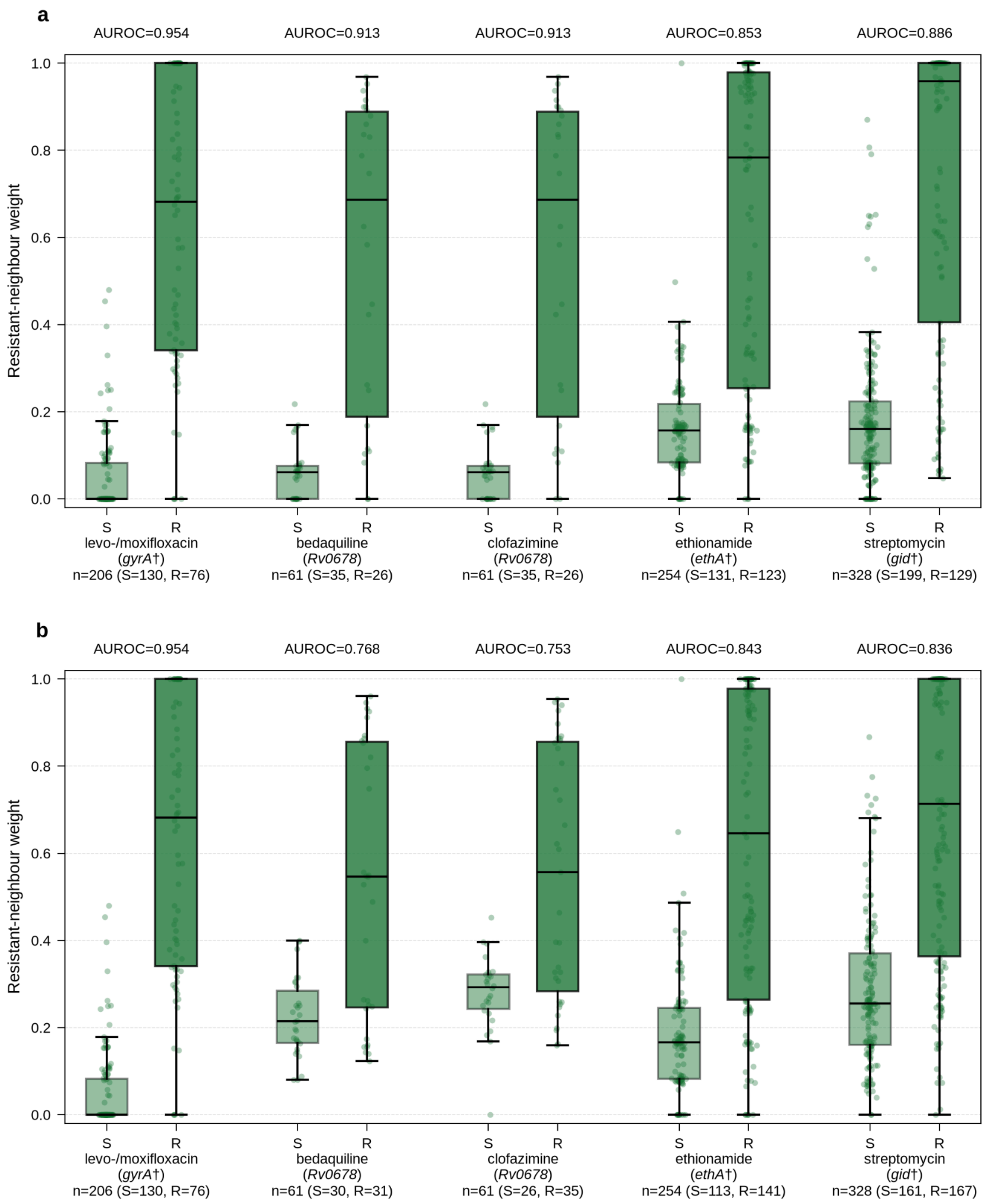


**Supplementary Data Fig. 9** | MicroDUET function-head neighbourhoods discriminate resistance for second-line antitubercular drugs. As Supplementary Data Fig. 8, for second-line drugs. †, some resistant isolates are called at a different gene from the one plotted levofloxacin 2 of 76, moxifloxacin 2 of 78, ethionamide 15 of 141, streptomycin 28 of 167.
